# Improved detection of evolutionary selection highlights potential bias from different sequencing strategies in complex genomic-regions

**DOI:** 10.1101/2021.09.28.462165

**Authors:** Tristan J. Hayeck, Yang Li, Timothy L. Mosbruger, Jonathan P Bradfield, Adam G. Gleason, George Damianos, Grace Tzun-Wen Shaw, Jamie L. Duke, Laura K. Conlin, Tychele N. Turner, Marcelo A. Fernández-Viña, Mahdi Sarmady, Dimitri S. Monos

## Abstract

Balancing selection occurs when multiple alleles are kept at elevated frequencies in equilibrium due to opposing evolutionary pressures. A new statistical method was developed to test for selection using efficient Bayesian techniques. Selection signals in three different data sets, generated with variable sequencing technologies, were compared: clinical trios, HLA NGS typed samples, and whole-genome long-read samples. Genome-wide, selection was observed across multiple gene families whose biological functions favor diversification, revealing established targets as well as 45 novel genes under selection. Using high-resolution HLA typing and long-read sequencing data, for the characterization of the MHC, revealed strong selection in expected peptide-binding domains as well as previously understudied intronic and intergenic regions of the MHC. Surprisingly, *SIRPA*, demonstrated dramatic selection signal, second only to the MHC in most settings. In conclusion, employing novel statistical approaches and improved sequencing technologies is critical to properly analyze complex genomic regions.

## Introduction

Balancing selection occurs when one or multiple sources of evolutionary pressures such as pleiotropy, overdominance, negative selection, and positive selection strike a balance to keep multiple competing alleles in equilibrium across a population. This is in contrast with negative selection, which purges alleles that are detrimental to fitness ^1–4^, and positive selection, which pushes advantageous alleles towards fixation ^5–7^. When balancing selection occurs, it not only results in increased polymorphism at the allele directly under evolutionary pressure, but surrounding variants on the same haplotypes will also rise in frequency, in a process known as hitchhiking (Figure 1). This leaves behind linkage disequilibrium (LD) blocks, regions that contain strong correlation among neighboring variants, and a higher local density of polymorphisms than would be expected from neutral genetic drift. Improved detection and understanding of balancing selection in the human genome can provide valuable insight into heritable diseases and our species’ adaptation to varying environmental exposures ^8,9^. Existing methods for identifying balancing selection look for enrichment of common alleles ^10,11^ or deviations from neutral drift ^12^, while others search for trans-species selective alleles ^9,13–16^. Testing for deviations from neutral drift may miss selective signals and testing for trans-species selective alleles predominantly captures only ancient signals that affect the fitness across multiple species. To address the shortcomings, we developed LD approximate Bayesian factor (LD-ABF), a new robust statistical method that directly investigate balancing selection by testing for both density of polymorphisms and strength of LD. Patterns of balancing selection were investigated using three distinct datasets derived from varying sequencing technologies. First, we scanned for selection genome-wide using phased high-quality SNP array and exome sequence data from 497 clinical samples (including 334 trios, Table 1). In order to investigate the major histocompatibility complex (MHC), a complex genomic region governing immunity known to be under various evolutionary pressures, in greater detail, we then specifically analyzed key human leukocyte antigen (HLA) genes using high-resolution Next Generation Sequencing (NGS) typing on thousands of unrelated haplotypes worldwide from the 17th International HLA and Immunogenetics Workshop (IHIW) ^17^. Finally, we validated our findings and identified complex signal artifacts using an independent set of high quality long-read whole genome sequencing (WGS) samples from the Human Pangenome Reference Consortium (https://humanpangenome.org).

**Figure 1.**
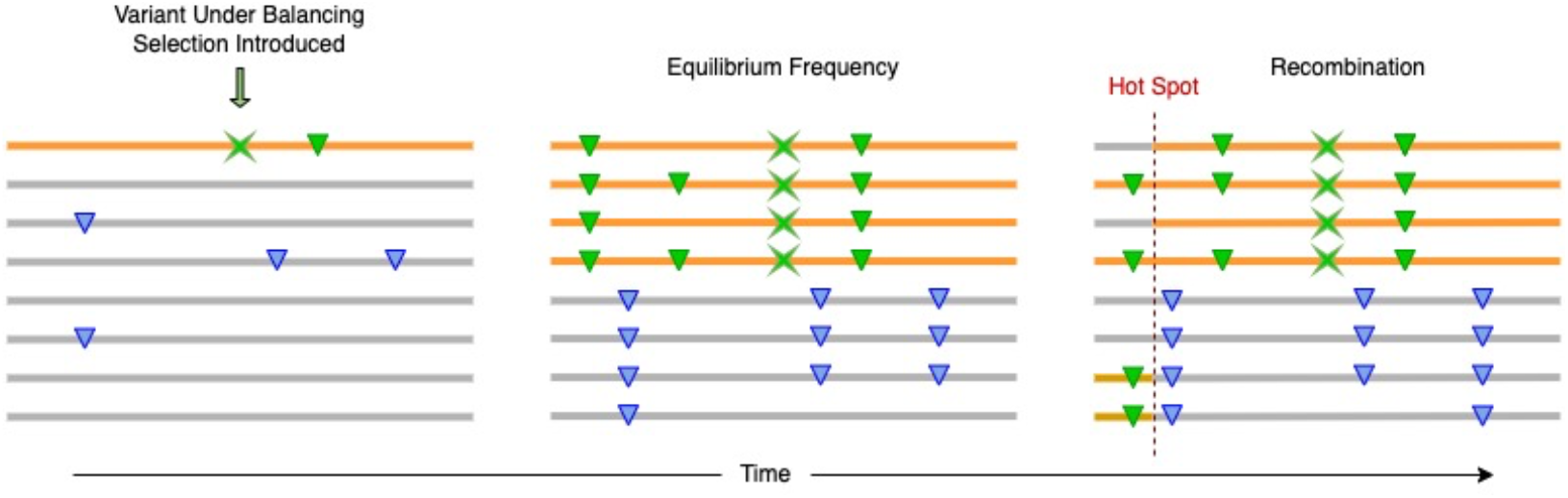
Evolutionary diagram depicting the progression of an allele under balancing selection. The green X denotes the variant under selection, green triangles are variants originating on the same haplotype denoted by an orange line as the balancing selection variant, and blue triangles occur on an alternate haplotype denoted by an orange line. In the first pane the variant is introduced on a single haplotype. Then after some time has passed evolutionary pressures favoring multiple alleles at the position of focus maintaining both haplotypes with and without the polymorphism, where hitchhiking effects are observed around the variant under balancing selection–inducing LD patterns. Recombination breaks the strong LD resulting in mosaics of the haplotypes, where strong hotspots will diffuse the LD effects of hitchhiking.

**Table 1.** Detailed counts for CHOP trios and individuals collected for analysis that include both SNP array data and whole exome sequence data.

| *Population* | *Individuals* | *Duo Probands* | *Trio Probands* | *Totals* |
| --- | --- | --- | --- | --- |
| **AFR** | 16 | 9 | 34 | 59 |
| **AMR** | 12 | 12 | 44 | 68 |
| **EAS** | 11 | 1 | 17 | 29 |
| **EUR** | 64 | 33 | 221 | 318 |
| **SAS** | 3 | 2 | 18 | 23 |
| **Totals** | 106 | 57 | 334 | 497 |

## Results

### Overview of LD-ABF Statistical Method

Approaches to assess balancing selection by quantifying local polymorphisms and LD patterns are complicated by both rare variants (resulting in sparse data) and instances of close or perfect LD among variants (resulting in quasi or fully separated data). We implemented a Bayesian logistic regression model using logF priors (the conjugate family for binomial logistic regression), which enabled us to utilize established data augmentation techniques to efficiently estimate posterior coefficients ^18–20^. Such application of logF priors, which are weakly informative priors that are also grounded in penalized regression methods, has been shown to be effective in settings of both sparse and fully separated data without making major assumptions ^18,21^. Then to test how well a SNP predicts its neighboring variants we derived an approximate Bayes factor (ABF)^22^. Finally, the log of the products of ABFs for every base in a set window (here 1Kb was used) is taken to derive a combined score that measures both the density of polymorphisms and degree of LD around the test SNP. Comparing against existing methods, evolutionary simulations showed that our novel method performed as well or better in almost all settings and appears most robust in picking up subtle signals of recent balancing selection (Supplemental Table 1 and Supplemental Figure 1). A more detailed account of the method can be found in the supplemental material and code is available online at https://github.com/tris-10/LD-ABF.

### Genome Wide Scan for Balancing Selection in Clinical Trios

First, we analyzed 497 clinical samples from the Children’s Hospital of Philadelphia with SNP array data and matching whole exome sequencing, including 334 trios (Table 1). These samples were phased using SHAPEIT2 and clustered into ancestral populations based on PCA using 1000 Genomes Project (1KGP)^23^ super populations^24,25^ (see Methods). LD-ABF was calculated genome wide for each population to determine where different balancing selection events occurred and in what populations (Figure 2A, Supplemental Figure 3-4). Although LD will dissipate further away from a selection event, there is some spread beyond the immediate window to neighboring regions. To identify unique selection events, when a local LD-ABF peak was identified, bases within a set neighborhood were excluded from additional LD-ABF peak determination. To be conservative in avoiding double counting peaks within long extended LD, the analysis was first performed using neighborhoods of 1 Mb around the highest local scores. A follow up analysis was then performed using 100 Kb neighborhoods to detect peaks at a finer granularity (Online Data). Within each super population, coordinates of the 100 highest peaks were used to identify candidate genes under balancing selection (Online data). Among these, 61 genes were shared across populations (Figure 2D), including key HLA genes. Furthermore, we investigated the top 10 peaks of each population in detail (Table 2).

**Figure 2.**
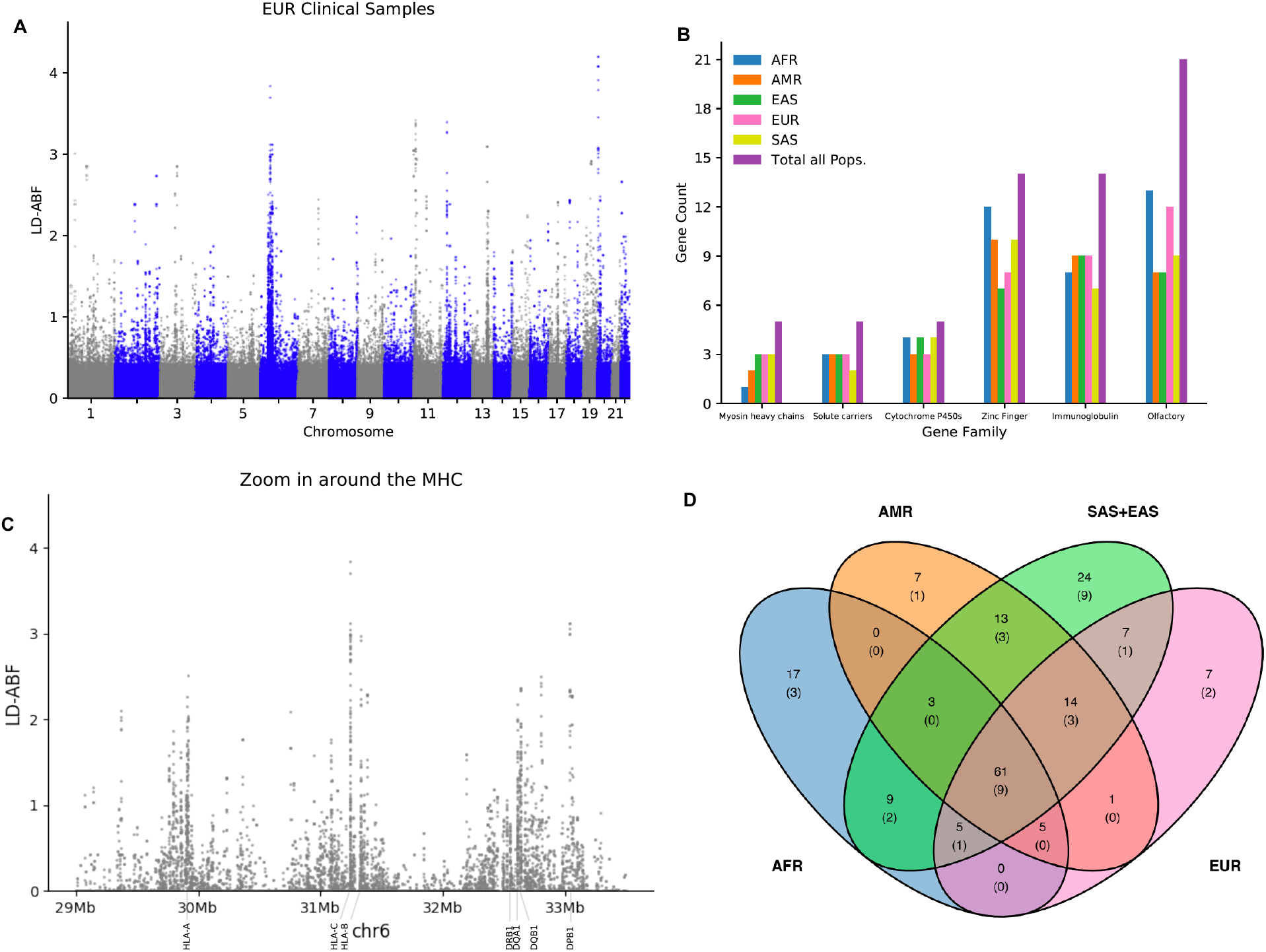
Genome wide scan for balancing selection in clinical samples and gene patterns. Clinical samples were clustered based on 1KGP superpopulations: African (AFR), American (AMR), East Asian (EAS), Southern Asian (SAS), and European (EUR). Genome wide scans were performed within population to detect balancing selection, here in A.) EUR genome wide with other populations shown in Supplemental Figure 3 and C) zoomed in plot across the MHC with class I and II HLA genes in the EUR clinical samples. Looking across the entire MHC, there appears to be several clusters of balancing selection signals centered around HLA genes. Three of these clusters (1. HLA-C, HLA-B; 2. HLA-DRB1, HLA-DQA1, HLA-DQB1; and 3. HLA-DPA1, HLA-DPB1) are separated by previously noted recombination hotspots^64–66^. Then restricting to the top 100 peaks, where LD-ABF scores in the immediate 100 Kb window around a peak are ignored to determine subsequent peaks, within each population is intersected with different B.) HGNC gene families to get gene counts and the D.) Venn diagram of unique and shared top 100 peak genes between populations with the two Asian populations combined with novel gene counts shown in parenthesis.

**Table 2.** Top 10 genome wide peaks in balancing selection signal in each clinical sample population. Peaks reported using 1Mb neighbor hoods with genic context and regional gene density. For instances where the exact peak position occurs at multiple variants within a region in perfect LD, the start and end positions are represented here and each individual variant can be found in the online data.

| Pop | Chr | Start | End | LD- ABF | Gene | Gene Category | # Genes within 100Kb | # Genes within 1Mb |
| --- | --- | --- | --- | --- | --- | --- | --- | --- |
| AFR | 11 | 5373251 | 5373251 | 0.75 | OR51B6 | Olfactory Receptor Family | 4 | 49 |
|  | 18 | 11609727 | 11610121 | 0.65 | SLC35G4 | Solute carrier family | 2 | 10 |
|  | 6 | 33037080 | 33037082 | 0.65 | HLA-DPA1 | Major Histocompatibility Complex, Class II | 3 | 47 |
|  | 20 | 1895889 | 1896100 | 0.64 | SIRPA | Signal regulatory protein | 2 | 14 |
|  | 1 | 158725194 | 158725194 | 0.63 | OR6K6 | Olfactory Receptor Family | 4 | 26 |
|  | 11 | 4790671 | 4790671 | 0.59 | OR51F1 | Olfactory Receptor Family | 2 | 38 |
|  | 13 | 93969248 | 93969473 | 0.52 | GPC6 | Glypican | 0 | 3 |
|  | 6 | 31379773 | 31379795 | 0.51 | MICA | Major Histocompatibility Complex, Class I | 3 | 81 |
|  | 22 | 22869123 | 22869218 | 0.49 | ZNF280A | Zinc Finger Protein | 4 | 12 |
|  | 11 | 7817852 | 7817959 | 0.47 | OR5P2 | Olfactory Receptor Family | 2 | 19 |
| AMR | 20 | 1895963 | 1895963 | 1.00 | SIRPA | Signal regulatory protein | 2 | 14 |
|  | 6 | 31237802 | 31237802 | 0.91 | HLA-C | Major Histocompatibility Complex, Class I | 2 | 73 |
|  | 11 | 5443887 | 5443887 | 0.73 | OR51B5,OR51Q1 | Olfactory Receptor Family | 5 | 48 |
|  | 11 | 7817852 | 7817856 | 0.71 | OR5P2 | Olfactory Receptor Family | 2 | 19 |
|  | 6 | 33037412 | 33037424 | 0.66 | HLA-DPA1 | Major Histocompatibility Complex, Class II | 3 | 47 |
|  | 3 | 75786737 | 75786737 | 0.57 | ZNF717 | Zinc Finger Protein | 2 | 8 |
|  | 19 | 41386420 | 41386420 | 0.57 | CYP2A7 | Cytochrome P450 proteins | 4 | 33 |
|  | 1 | 24201448 | 24201448 | 0.57 | CNR2 | Cannabinoid receptor | 3 | 24 |
|  | 22 | 22869123 | 22869218 | 0.56 | ZNF280A | Zinc Finger Protein | 4 | 12 |
|  | 13 | 93969248 | 93969473 | 0.55 | GPC6 | Glypican | 0 | 3 |
| EAS | 20 | 1895889 | 1896060 | 0.42 | SIRPA | Signal regulatory protein | 2 | 14 |
|  | 6 | 33037412 | 33037412 | 0.37 | HLA-DPA1 | Major Histocompatibility Complex, Class II | 3 | 47 |
|  | 6 | 31237802 | 31237802 | 0.30 | HLA-C | Major Histocompatibility Complex, Class I | 2 | 73 |
|  | 11 | 5443887 | 5443887 | 0.27 | OR51B5,OR51Q1 | Olfactory Receptor Family | 5 | 48 |
|  | 1 | 248525328 | 248525330 | 0.26 | OR2T4 | Olfactory Receptor Family | 5 | 37 |
|  | 1 | 89652071 | 89652090 | 0.24 | GBP4 | Guanylate-binding proteins | 2 | 16 |
|  | 22 | 22869123 | 22869218 | 0.24 | ZNF280A | Zinc Finger Protein | 4 | 12 |
|  | 14 | 105418234 | 105418235 | 0.23 | AHNAK2 | PDZ domain containing | 3 | 27 |
|  | 1 | 24201448 | 24201448 | 0.23 | CNR2 | Cannabinoid receptor | 3 | 24 |
|  | 6 | 159654994 | 159654994 | 0.22 | FNDC1 | fibronectin type III domain containing 1 | 1 | 13 |
| EUR | 20 | 1895990 | 1895990 | 4.20 | SIRPA | Signal regulatory protein | 2 | 14 |
|  | 6 | 31237876 | 31237876 | 3.84 | HLA-C | Major Histocompatibility Complex, Class I | 2 | 73 |
|  | 11 | 5373242 | 5373242 | 3.43 | OR51B6 | Olfactory Receptor Family | 4 | 49 |
|  | 12 | 11244390 | 11244390 | 3.40 | PRH1,PRH1-PRR4,<br>PRH1-TAS2R14,TAS2R43 | Heterogeneous family of proline-rich salivary glycoproteins,<br>taste receptor | 3 | 31 |
|  | 6 | 33037419 | 33037424 | 3.13 | HLA-DPA1 | Major Histocompatibility Complex, Class II | 3 | 47 |
|  | 13 | 93969248 | 93969473 | 3.10 | GPC6 | Glypican | 0 | 3 |
|  | 11 | 244106 | 244167 | 3.05 | PSMD13 | 26S Proteasome, a multicatalytic proteinase | 9 | 36 |
|  | 1 | 24201448 | 24201448 | 3.02 | CNR2 | Cannabinoid receptor | 3 | 24 |
|  | 11 | 7817852 | 7817959 | 2.94 | OR5P2 | Olfactory Receptor Family | 2 | 19 |
|  | 19 | 41386420 | 41386420 | 2.92 | CYP2A7 | Cytochrome P450 proteins | 4 | 33 |
| SAS | 6 | 33037412 | 33037424 | 0.33 | HLA-DPA1 | Major Histocompatibility Complex, Class II | 3 | 47 |
|  | 20 | 1895889 | 1895990 | 0.30 | SIRPA | Signal regulatory protein | 2 | 14 |
|  | 12 | 11244378 | 11244390 | 0.24 | PRH1,PRH1-PRR4,<br>PRH1-TAS2R14,TAS2R43 | Heterogeneous family of proline-rich salivary glycoproteins,<br>taste receptor | 3 | 31 |
|  | 1 | 248525328 | 248525330 | 0.21 | OR2T4 | Olfactory Receptor Family | 5 | 37 |
|  | 11 | 5443887 | 5443887 | 0.20 | OR51B5,OR51Q1 | Olfactory Receptor Family | 5 | 48 |
|  | 1 | 24201448 | 24201448 | 0.19 | CNR2 | Cannabinoid receptor | 3 | 24 |
|  | 19 | 41386136 | 41386136 | 0.18 | CYP2A7 | Cytochrome P450 proteins | 4 | 33 |
|  | 2 | 234622061 | 234622110 | 0.18 | UGT1A10,UGT1A5,<br>UGT1A6,UGT1A7,<br>UGT1A8,UGT1A9 | UDP-glucuronosyltransferase | 9 | 22 |
|  | 22 | 22869123 | 22869218 | 0.17 | ZNF280A | Zinc Finger Protein | 4 | 12 |
|  | 13 | 93969248 | 93969473 | 0.17 | GPC6 | Glypican | 0 | 3 |

The top peak for the AFR population is in *OR51B6* of the olfactory receptor (OR) gene cluster; for SAS, the top peak appears in *HLA-DPA1*, a MHC class II gene; and for AMR, EUR, and EAS populations, the top peak is in *SIRPA*, which encodes for a signal regulatory protein of the immunoglobulin superfamily. In fact, peaks in *SIRPA* rank among the top 4 for each population. The second strongest selection signal in the AFR samples is in *SLC35G4*, which encodes for a putative solute carrier. This strong selection signal in *SLC35G4* is a novel one. Among all top 100 peaks across populations, a total of 45 novel genes (online data, 34 genes using 100Kb peak neighborhood Figure 2D, 37 with 1Mb peak neighborhood) were tagged by signals of balancing selection (Table 2 and online data), including 9 shared between all populations (Figure 2D): *COL5A1, HCG20, OR1S1, OR2T4, QRICH2, SLC35G4, SNHG14, SNRPN, TRMT9B*.

As expected, several other top peaks are in HLA genes. In fact, peaks in *HLA-A, -C*, and *-DPA1* are shared among the top 100 peaks across all populations. Their relative rankings, however, vary from population to population. In the largest populations, EUR and AMR, the highest HLA peak is found in *-C*, while for AFR, EAS, and SAS, the highest HLA peak is found in *-DPA1*. In total, 18 HLA and other immunoglobulin superfamily genes are marked by top 100 LD-ABF peaks across all populations (Figure 2B and Supplemental Table 4). Immune related and cell surface receptor signaling genes are expected candidates for balancing or positive selection as their functionality is often directly tied to environmental interactions. Consistent with this, we also detected LD-ABF peaks across 22 OR genes and several taste receptor genes (Figure 2B and online data). In addition, peaks were also seen across members of several other gene families ^26^, including zinc fingers (ZF) (14), cytochromes (6), solute carriers (4), and myosin heavy chains (4) (Figure 2B).

Bases scoring in the top 0.1% LD-ABF genome wide were then intersected with known GWAS catalog significant SNPs ^27^ to find overlap between strong signals of selection and known disease associated variants (Table 3 and Supplemental Table 2). Using 0.1% coincides with a more restrictive threshold than the cut off for top 100 peaks while still allowing for consideration of multiple variants of interest within the same peak. Many of the SNPs overlapping high LD-ABF scores were found to be associated with blood and immune related traits. Among these, the strongest signal for EAS was at rs17855611 in *SIRPA* associated with blood protein levels, and for SAS, at rs1126506 in *HLA-DPA1* associated with anti-rubella IgG levels. In contrast, the strongest signals in AFR, AMR, and EUR were seen in *OR51B6*, which corresponds to rs5006884 with known association to fetal hemoglobin (HbF) levels in sickle cell anemia, a classical example of balancing selection driven disease ^28^. This SNP lies upstream of the β-globin locus control region and is in close proximity to several candidate enhancers of *HBG2* ^29^, which codes for the gamma-2 subunit of HbF. ClinVar SNPs ^30^ were also investigated, showing possible selection in *CYP2D6* and *OPRM1* related to drug responses, and in *IRF5* and *HAO*, associated with systemic lupus erythematosus and calcium oxalate urolithiasis respectively (see Methods and Supplemental Table 5).

**Table 3.**
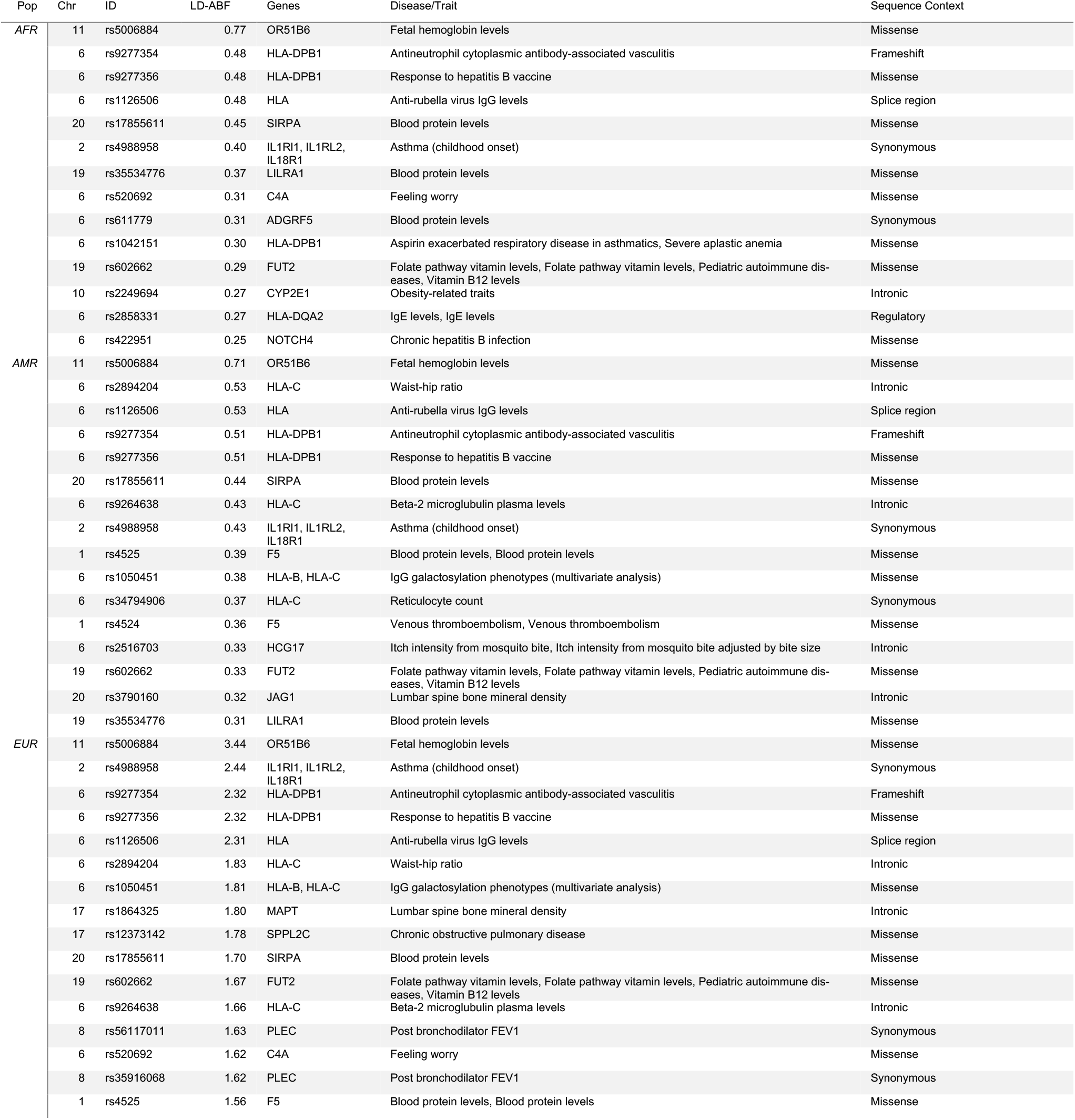
Top balancing selection signals in clinical samples at GWAS significant SNPs. SNPs that are both found to be significantly associated with a phenotype in the GWAS catalog and also have a strong selection signal in the top 0.1%. The results for clinical samples in the EUR, AFR, and AMR populations are here with the EAS and SAS populations continued in Supplemental Table 2.

#### Detailed look at HLA Genes Using High Quality Typing

Diversity in HLA genes have long been recognized as key examples of balancing selection^31–33^. Moreover, even though the MHC accounts for only 0.16% of the genome, 39% of all GWAS SNPs that overlapped top LD-ABF scores occurred within the MHC. Despite these observations and its profound importance to the fields of immunology, immunogenetics, and evolutionary biology, detailed follow up and characterization of the MHC and its HLA genes has been limited. Fortunately, due to the vital importance of HLA matching for avoiding rejection and graft versus host disease in organ and stem cell transplants, detailed typing of selective HLA genes is routinely performed in the clinical setting ^34–36^.Taking advantage of this, we utilized high-resolution HLA typing data from the IHIW to take a closer look at balancing selection across these genes. This dataset consists of over 3,500 samples, each providing 2 alleles per HLA gene typed at 4 field resolution and represents a diverse set of world populations (see Methods).

Strikingly, the strongest LD-ABF signals were consistently observed in -*DQA1*, -*DQB1*, and *DRB1* across all IHIW populations (Figure 3 and Supplemental Figure 11-13). This is in contrast to scans of the clinical samples, where either *-C* or *-DPA1* were the top hits across the MHC depending on the population. Furthermore, within each HLA gene, consistent patterns of balancing selection were observed across all populations, including strong signals in the intronic regions (Supplemental Figure 7-13). Not surprisingly, these regions with the highest LD-ABF scores corresponds to regions with the highest concentration of GWAS trait associated SNPs. A review of SNPs overlapping top LD-ABF scores revealed associations with traits like red blood cell count, leukemia, autism, schizophrenia, and asthma (Supplemental Table 3). The sequence context of the majority of these SNPs was either intronic or missense, which is expected in the context of balancing selection; as opposed to nonsense or frameshift SNPs, which would be expected in settings of purifying selection. Looking over the exons of HLAs, the highest LD-ABF signals for both *-DQA1* and *-DQB1* were found in exon 2, which encode for extracellular domains key to peptide presentation. Diversity in the peptide-binding pocket ensures effective immune recognition of a wide range of foreign pathogens, in tune with mechanisms driving balancing selection.

**Figure 3.**
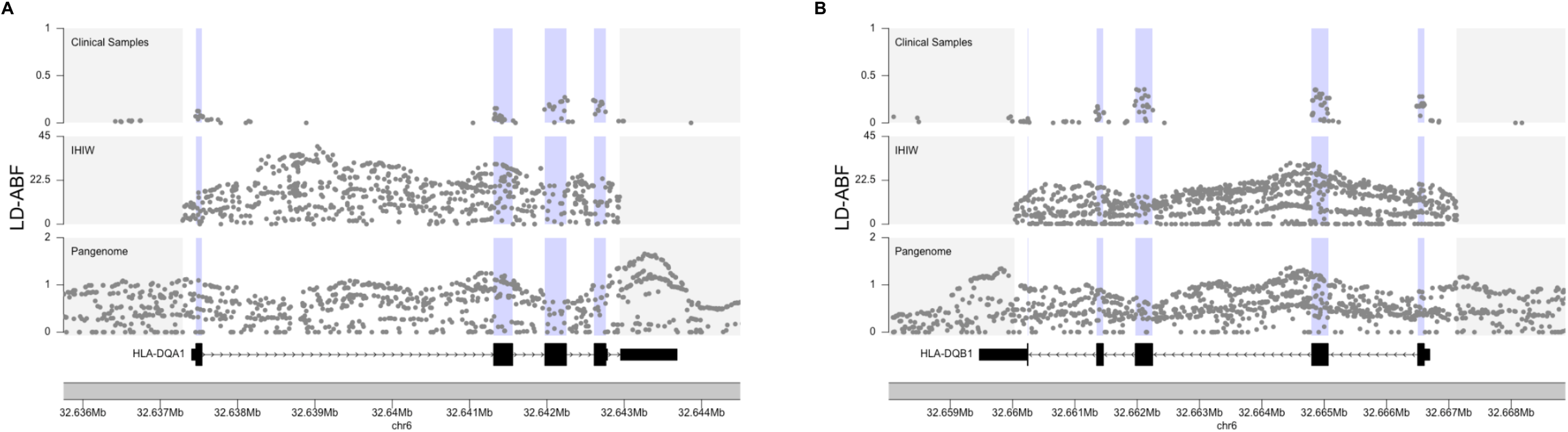
Balancing selection in HLA-DQA1 and DQB1 comparing the clinical samples, 17^th^ IHIW, and Pangenome. LD-ABF scores over A) DQA1 and B) DQB1 from independent samples of African ancestry are compared. Exonic regions are highlighted in purple. The relative magnitude of the LD-ABF signals reflects the sample size of the population as any standard test statistic would.

### Validation with Long-read Pangenome Samples

To further validate and reconcile findings, LD-ABF testing on whole genome HiFi PacBio sequencing data gathered by the Pangenome Consortium was performed. These high quality long-read samples are expected to help remove artifacts introduced by inaccurate assembly and alignment of other platforms. This is especially applicable for genomic regions of high homology and complexity that are difficult or impossible to properly align and map when using short-read sequencing, including the MHC. Although these samples offer superior sequencing quality, the largest population consists of just 23 African samples (Supplemental Figure 14); so, they are presented here predominantly for selective verification and not as part of the broader analysis. The other Pangenome populations were too small to perform statistical inference.

Revealingly, with the African Pangenome samples, signals at *SIRPB1* seen in the clinical samples were absent (Supplemental Figure 5), indicating that they were likely artifacts of inaccurate sequence mapping. In contrast, the strong signals in the MHC and *SIRPA* were again demonstrated, even with the more restrictive segmental duplication filter applied (Figure 4). The magnitude of the *SIRPA* signal is second only to the MHC in the Pangenome data, confirming strong balancing selection. Beyond the MHC and *SIRPA*, the top 100 peaks in the Pangenome samples (online data) included *OR51B5, MYO3A*, and *OR6J1*, which were also found to be top hits for clinical samples. Additionally, when removing the segmental duplication filter, two more genes, *LILRA6* and *FLG*, overlapped. While *LILRA6* appears to be another balancing selection candidate of interest, we caution any inferences to be made on *FLG* as its signal appears borderline.

**Figure 4.**
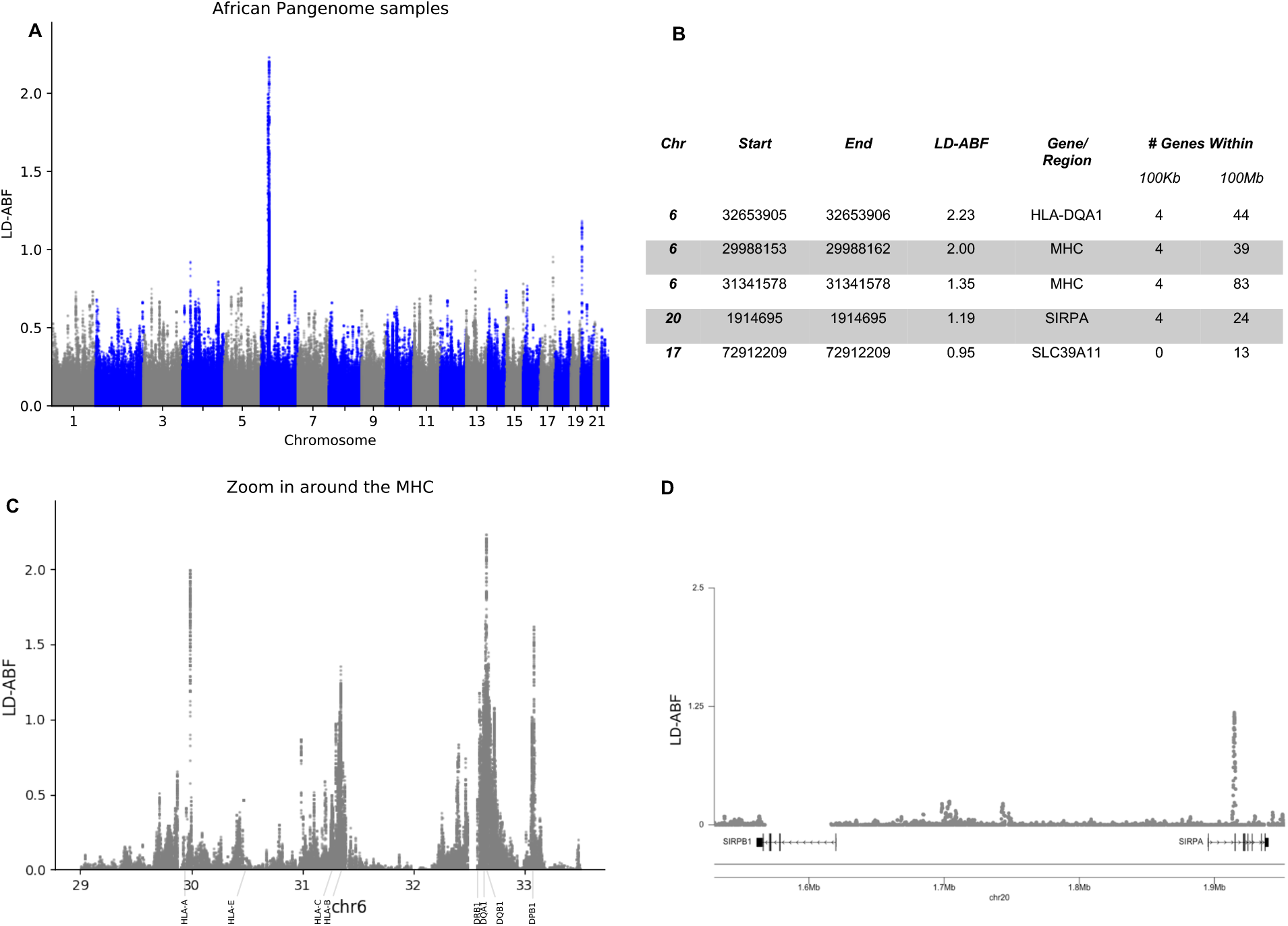
Signals of balancing selection detected in the Pangenome samples. LD-ABF scores calculated from long-read HiFi PacBio data are shown A) genome wide and with a B) table detailing the top 5 LD-ABF peaks C) zoom in around the MHC D) zoom in around the SIRP genes.

Signals in HLA genes from African populations were compared across datasets. As the scale of LD-ABF signal is a function of sample size, for this comparison, we focus on the relative peaks and shapes of the distributions as opposed to the absolute LD-ABF scores. Since the data for the clinical samples are limited by the exome sequencing and variants on the SNP arrays, it became clear how incomplete the data were as compared to the IHIW and the Pangenome (Figure 3 and Supplemental Figure 15). The patterns of LD-ABF from the IHIW samples largely matched those of the Pangenome samples, with the exception of a problematic subregion within the *HLA-DRB1* (Supplemental Figure 15 and Supplemental Figure 13). A dramatic peak centered on intron 5 of -*DRB1* seen in the IHIW dataset was completely absent in the Pangenome analysis. This portion of DRB1 is known to have structural variation and repeat elements, hindering accurate mapping of shorter sequencing reads, and therefore likely causes artifactual LD in IHIW but not the Pangenome (see Methods). The Pangenome, and long read sequencing in general, offers an invaluable resource for reconciling such artifacts while also providing dramatic replication of surprisingly strong signals like that seen in *SIRPA*.

## Discussion

LD-ABF improves detection of evolutionary selective pressures by measuring both the strength of LD and the density of variation. Here, we analyzed three independent datasets representing different sequencing technologies, each with unique advantages and limitations. The comparison revealed the significant impact of sequencing strategies in identifying patterns of selection that likely applies to any such study of evolutionary pressures.

The MHC is a genomic region of particular interest both from a medical perspective and in terms of understanding evolutionary pressures. Studies have linked over 700 diseases and traits to the MHC, more than to any other genomic region of comparable size ^37,38^. In fact, SNPs within the MHC represent nearly 2% of all GWAS catalog associations genome wide ^27^. Much work has also been done looking at the MHC as a key example of balancing selection ^31–33^, with an emphasis for greater selection in class I genes ^39^. In general agreement with prior studies, we also saw some of the strongest LD-ABF scores genome wide within the MHC. However, there is a limitation of using SNP array and exome data alone, as it is inherently restricted to detecting evolutionary selection only on the variants covered by the platforms. In this study, we utilized thousands of samples reported by the 17^th^ IHIW to better characterize key HLA genes within the MHC. With this improved resolution, interestingly we saw the strongest signals in *HLA-DQA1*, -*DQB1*, and -*DRB1* across all populations. Supporting the IHIW results, the African Pangenome samples also showed the strongest signals in the DQ region. Looking at -*DQA1* and -*DQB1* in more detail using both datasets, the strongest exonic signals appeared in exon 2 for both genes, which codes for α1 and β1 subunits respectively of the peptide-binding domain. Interestingly, when examining class I HLA genes, this was not always the case. For both class I and class II genes, exons coding for the peptide-binding domains are known to be the most diverse, consistent with selection in the presence of varying pathogen exposures. Importantly, while the peptide-binding domain is derived from a single gene for class I, it is formed by the dimerization of α and β chains encoded by separate genes in class II. Each haplotype’s α chain is capable of dimerizing with not only its β chain in *cis*, but also with the β chain of the opposite haplotype. While the α chain of DR is nearly monomorphic, both α and β chains are highly polymorphic for DQ and DP. Critically, only certain combinations of α and β alleles are frequently observed, which are reinforced by known LD between *-DQA* and *-DQB* and *-DPA* and *-DPB*. These same selective restraints likely contribute to the strong LD within the DQ genes centered around exon 2. In contrast, *DPB1* appears to demonstrate a different LD pattern from other class II genes, with stronger selection signal in the intronic regions as compared to exonic regions dominated by only a few alleles (Figure 3 and Supplemental Figure 15). There is a known recombination event in the DP region ^40–42^ and the -*DPA1* and -*DPB1* genes overlap with opposite orientations. Additionally, there are multiple functional elements within this overlap region, including multiple eQTLs, ^43,44^ two promoters (one for each gene), and a processed pseudogene of the ribosomal protein L32 ^45^, further constraining this portion of sequence. It is noteworthy that correspondingly, we saw a distinct dip in the LD-ABF (Supplemental Figure 17) over this region.

The patterns of LD-ABF in HLA genes are consistent across different populations based on the IHIW data (Supplemental Figure 7-13); however, they are variable across different populations based on the SNP array and exome data of the clinical samples (Table 2 and Supplemental Figure 4). This strongly suggests that the previously observed variability in balancing selection between populations, at least in part, is due to sparce data that’s inherent of SNP arrays and even exome sequence data (Figure 3 and Supplemental Figure 15). Beyond highlighting the strong balancing selection signals in class II HLAs, the IHIW and Pangenome data also revealed very strong signals in intronic and intergenic regions of the MHC (Figure 4, Figure 3, Supplemental Figure 15 and Online Data), which have not been extensively analyzed by previous studies. Many GWAS disease associated SNPs fall within these noncoding regions; our analysis here begins to offer some clues regarding the evolutionary forces that contributed to these polymorphisms. Although the clinical samples also showed strong signals across HLA genes, it alone would have missed much of these interesting intricacies due to the sparseness of the data, especially over introns and intergenic regions. Furthermore, the consistent patterns of balancing selection in the HLA genes across different populations in the IHIW data (Supplemental Figure 7 Supplemental Figure 13) hints at possible convergent evolution, which have previously been noted in the HLAs^17,46^. Future work would look to distinguish between genetic similarity arising from ancestral adaptation being passed over generations versus convergence of different haplotypic lineages driven by similar selective pressures resulting in consistent genetic character.

In the clinical samples, *SIRPA* had one of the strongest, if not the strongest, balancing selection signals genome wide. The signal was replicated in the Pangenome, a completely independent sample set (Figure 4). Tennessen et al also observed selection around *SIRPA*, though they did not identify nearly as strong of a signal–likely due to a combination of different sequencing platforms, sample sources, and methodology. SIRPα acts as an inhibitory receptor for CD47 and is a key component of the “do-not-eat-me” signaling pathway and may have implications in transplantations ^47^. Similar to the HLA genes, the strongest signal appears in sequences coding for the extracellular domain of SIRPα^48^. Interestingly, although this outward facing domain of SIRPα is analogous to the antigen-binding domains of HLAs and immunoglobulins ^49^, structural analysis showed that unlike variation in the complementary determining regions of those proteins, most polymorphisms in *SIRPA* do not affect CD47 binding ^50^. Instead, they cluster away from the CD47 binding footprint, and are thought to be selected to minimize pathogen binding and manipulation of the “do-not-eat-me” signal ^50^.

Beyond HLAs and *SIRPA*, several other notable genes and gene families were identified by top LD-ABF peaks across all populations (Table 2). OR genes formed the largest gene family under balancing selection. Notably, both HLAs and ORs are thought to have diversified through gene duplications and consequently both families reside in regions of high gene density. These observations, along with the high homology among members of HLAs, ORs, and other gene families identified by our method suggests that balancing selection and gene duplications are often the result of similar evolutionary pressures. Similarly, TAS2R genes, encoding bitter taste G-protein coupled receptors, also form a cluster and have been found to be under selective pressure ^51^. Although technically neither an OR nor a taste receptor gene, *CNR2*, coding for a cannabinoid receptor, was also identified in the top 10 peaks of several populations. It is known to have associations with psychoactive and anti-inflammatory responses ^52,53^. Following ORs and the immunoglobulin superfamily, ZFs form the third largest gene family under selection and includes 2 genes identified in top 10 peaks: *ZNF280A* and *ZNF717*. Since ZFs function as binding molecules, with DNA and RNA among their targets, it comes as no surprise that polymorphism dictating binding specificity were found to be under balancing selection. Additionally, several cytochrome P450 genes were also identified. These enzymes catalyze many reactions in drug metabolism and lipid synthesis ^54^; their polymorphisms have been extensively studied and are of vital importance in pharmacology. Furthermore, *GPC6* was found within top 10 peaks in four of five populations demonstrating dense LD over several Mbs (Supplemental Figure 6), similar to those seen around HLAs. *GPC6* is associated with bone density ^55^ and omodysplasia ^53,56^, a rare skeletal dysplasia characterized by severe limb shortening^57^. *GBP4*, with a top 10 peak in EAS and top 100 peaks across every other population, is an IFN-inducible GTPase of the guanylate binding protein family, whose members has emerged as key orchestrators of inflammation in anti-bacterial immunity, metabolic disorders, and cancer ^58^.

Looking at the top 100 peaks across every population (1Mb and 100kb neighborhoods), we identified a total of 45 novel genes. Although these specific genes were not previously described to be under selection, related genes or their gene families have been found by previous studies. Among these, *SLC35G4* had the second strongest signal in the AFR clinical samples (and top 100 in all other populations) and was corroborated by the Pangenome analysis (Supplemental Figure 16). *SLC35G4* belongs to the solute carrier family of genes, which has been found to be under selection^7,9,15,59^. Although minimally studied to date, SLC35G4 has recently been described as a potential neoantigen in prostate cancer^60^.

Several limitations of this work leave room for future investigation. When evaluating the representativeness of our datasets, it must be noted that the clinical samples correspond to individuals that have come into the children’s hospital for various clinical assessment and not specifically curated for the specific study of evolutionary selection. Depending on the dataset, there were limitations directly noted in terms of sequencing quality and/or representation of certain populations. The current analysis focuses on LD within a 1Kb window and does not test for long-range LD. To these ends, long-read sequencing will become increasingly important^61^ as we try to decipher the complexity of the MHC and other regions of genome with high homology or extensive LD.

Our results demonstrated that orders of magnitude smaller set of high-quality long-read sequencing data has the potential to more effectively characterizing genetic variation than larger sets of sequencing data from other platforms. Potentially, a combination of high-quality sequencing data and an optimal set of samples, would offer the most cost-effective way of performing such studies while providing thorough characterization of complex genomic regions. In addition, improved mapping and alignment techniques, like the use of population reference graphs^62^, will further facilitate genetic characterization of different human populations^63^. This, coupled with methodological advances like LD-ABF, will enable the better understanding of evolutionary pressures and their impacts on genomic functionality as well as the interrelationships between pathogens and corresponding diseases.

## Acknowledgements

We would like to thank both the participants in 17^th^ IHIW, including the patients and donors who volunteered to have their samples collected and analyzed for research purposes, and the teams across the world that performed the typing and collection of samples. Thank you to Joseph Antonelli for his insightful comments about the statistical approach. We would also like to thank Steven Pastor for his helpful feedback about the patterns observed over DRB1.

## Online Resources

IHIW http://17ihiw.org/17th-ihiw-ngs-hla-data/

IMGT https://www.ebi.ac.uk/ipd/imgt/hla/

NCBI gene database https://www.ncbi.nlm.nih.gov/gene/

GeneCards www.genecards.org

Pangenome https://s3-us-west-2.amazonaws.com/human-pangenomics/index.html?pre-fix=working/HPRC/HG01361/assemblies/

HUGO Gene Name Committee: https://www.genenames.org/data/genegroup/#!/group/589

Git repository with code and online data: https://github.com/tris-10/LD-ABF

In addition to the code, data files can be downloaded online data (github Readme.md section Download LD-ABF supplemental files):

- CHOP Trios: Genome Wide LD-ABF test statistics and peaks detailed for all included populations in Hg19
- All 17^th^ IHIW: HLA LD-ABF test statistics for all included populations, tab delimited sequence data generated from 17^th^ IHIW and IMGT 3.25 with lifted over alignments to Hg19 performed. Plots across all genes for all included populations.
- Pangenome Freeze 1 African samples: LD-ABF test statistics and variant calling vcfs in Hg38 for samples.

## Notes

http://17ihiw.org/17th-ihiw-ngs-hla-data/

https://www.ebi.ac.uk/ipd/imgt/hla/

https://www.ncbi.nlm.nih.gov/gene/

https://www.proteinatlas.org

https://s3-us-west-2.amazonaws.com/human-pangenomics/index.html?prefix=working/HPRC/HG01361/assemblies/

### Competing Interest Statement

The authors have declared no competing interest.

### Summary of Updates

Updated draft with full Genome Wide scan of Pangenome and filtering of additional outliers from clinical samples.

