## Supplemental Materials for "Improved detection of evolutionary selection highlights potential bias from different sequencing strategies in complex genomic-regions"

**Methods:**

**Children's Hospital of Philadelphia Clinical Samples**

In the clinical samples, 834 samples underwent quality control and outlier filtering, leaving 497 with SNP array data and matching whole exome sequencing, including 334 trios (Table 1). These remaining samples were phased using SHAPEIT2 prior to LD-ABF analysis [1,2]. Since signals of selection can often be obscured or confounded by demographic shifts across populations, inference on each sample's ancestry was completed to facilitate within-population analysis. Using the first 10 principle components (PCs) calculated from the SNP array data, K-nearest neighbors clustering algorithm was run to group samples by their best matching 1000 Genomes Project (1KGP) [3] super population—Africa (AFR), East Asia (EAS), Europe (EUR), South Asia (SAS), or the Americas (AMR) (Supplemental Figure 2). Sixteen outliers whose PC positions are more than six standard deviations away from the mean of any ancestral group were removed [4,5]. Such inference is expected to have limitations since the samples were not collected prospectively with ancestry or ethnicity assessments.

For samples to be included they needed to have both SNP array and whole exome sequence data, including 573 probands from trios and a total of 834 individuals. For the SNP array data, 832,381 SNPs common to the 3 SNP arrays were extracted. SNPs were then removed if they had genotyping call rate < 0.95, minor alleles frequency < 0.01. Individuals were removed if they had individual missing genotypes rates > 0.05. For the whole exome sequence data, within each family indels were separated from SNPs. Indels are excluded if QD < 2 or FS > 200 or ReadPosRankSum < -20. SNPs are excluded if QD < 2 or FS > 60 or MQ < 40 or MQRankSum < -12.5 or ReadPosRankSum < -8. Also within each family, genotypes variants were excluded if any individual had a variant with DP < 5 or GQ < 10. Vcf files were then merged across families and missing genotypes were assumed to be reference. Monomorphic, multi-allelic and variants with Mendel error rates > 0.01 were removed. The last 4 variants on chromosome 1 and chromosome 3 were also removed to finish the phasing.

Several transmission filters were incorporated leveraging family relatedness because certain variants appeared to be incorrectly called due to homologous sequence stretches. Some variants in these regions, of which appear to be paralogs, where heterozygosity was likely inaccurately called due to mapping problems demonstrated unrealistic transmission frequencies. Two tests were implemented to remove individuals that were excessively heterozygous. Relative to filtering out repeat masker regions entirely, this gives another way to remove potential artifacts by leveraging family data without having to remove close to 20% of variants. In settings where over 95% of families, either trios or duos, consisted of entirely heterozygous individuals those variants were filtered out. Then looking at complete trios, if both parents are heterozygous at an allele, the transmission of either homozygous variant is expected to be 25%. So, looking at each trio where both parents are heterozygous at a variant, a binomial test with a p-value threshold of 0.005 is constructed so the probability of success (ie seeing a homozygous proband) is p=25% and for the number of observations, n, is equal to the number of families with heterozygous parents; variants that do not pass the threshold are then filtered out. Sometimes the reference allele was not the major allele, ie the major and minor allele were flipped, in which case if the minor allele occurred more than 95% it was removed, this is the same as a 5% MAF threshold.

For both platforms, phasing was done using SHAPEIT2 [2] and then the cross platform samples were merged maximizing overlapping alternate allele matches. Several filters included removing regions that fell in the encode black list regions https://github.com/Boyle-Lab/Blacklist/ [6] low complexity repeat regions (LCR): https://raw.githubusercontent.com/lh3/varcmp/master/scripts/LCR-hs37d5.bed.gz, removing centromeres (acen) and telomers (gvar) UC genome browser and taking http://hgdownload.cse.ucsc.edu/goldenPath/hg19/database/cytoBand.txt.gz and any remaining indels. When running LD-ABF the within population variants were restricted to MAF > 0.05.

Top 100 peaks for each population are reported online. To be conservative in avoiding double counting peaks within long extended LD, the analysis was first performed using neighborhoods of 1 megabase (Mb) around the highest local scores. A follow up analysis was then performed using 100 kilobase (Kb) neighborhoods to detect peaks at a finer granularity (Online Data). Further analysis was doing looking at ClinVar variants. Since our test largely focuses on patterns of LD impacted by common variation as opposed to rare variation with strong deleterious effects under negative/purifying selection, it was largely expected that few top LD-ABF scores would correspond to ClinVar variant SNPs [7]. In fact, no overlap was detected between the top 0.1% LD-ABF scores and ClinVar variant SNPs. However, when we relaxed the threshold to include peaks in the top 1%, intersections with several CinVar variants related to drug response were found within *CYP2D6* and *OPRM1*. Additional overlaps were also found in *IRF5* and *HAO1*, in association with risks for systemic lupus erythematosus and calcium oxalate urolithiasis respectively (Supplemental Table 5).

**17th IHIW and IMGT**

Samples were taken from the 17th IHIW, using reported high resolution allele frequencies characterized by next generation sequencing in unrelated populations (17th IHIW Table 1). This dataset consists of over 3,500 samples, each providing 2 alleles per HLA gene typed at 4 field resolution and represents a diverse set of world populations: European Americans, African Americans, USA Hispanics, Spanish, Mexican, Italian, Greek, Asian Pacific Islanders, Thai, Indian, Arab, and Europeans (taken from the 17th IHIW Table 1). Since the samples reported include allele frequencies using classic HLA nomenclature, to perform analysis the data required matching on consensus sequencing then lifting over to reference. The observed alleles in 17th IHIW were matched with their established sequences, as described in the interational ImMunoGeneTics (IMGT) HLA database version 3.25.0, which is the version that most directly corresponds to the 17th IHIW and lifted over to Hg19. Indels, short tandem repeats (STRs), and missing variants were ignored for this analysis. In the 17th IHIW dataset, alleles ending with 'SG' in their name refer to short tandem repeat (STR) allele ambiguity groups; when encountering such alleles, we have removed the suffix to enable matching with a corresponding and representative IMGT allele. If an allele reported in 17th IHIW did not match up with a fully sequenced HLA allele in IMGT 3.25.0 then it is omitted. This typically only occurred with rare alleles, where all but DPB1*01:01:01 had allele frequencies below 5%. Low frequency alleles are expected to have less of an impact on the analysis than higher frequency alleles since LD is typically less strong for rare alleles. Genes without genomic alignment file for IMGT 3.25.0 were also omitted. Alleles were 4 field typed except where amplicons do not extend the full length of the gene where ambiguities are noted by the 17th IHIW (http://17ihiw.org/wp-content/uploads/2018/10/Readme-Unrelated-HLA-allele-and-haplotypes-FQ-tables_072318.pdf).

**Pangenome Samples**

Freeze 1 version 2 assembly data was downloaded from the Human Pangenome Reference Consortium (HPRC) repository. The assemblies were aligned to hg38 chromosome 6 using minimap2 (v2.21) in asm20 mode. All contigs with a total alignment length exceeding 500K were retained for variant calling. Filtered contigs were processed with Dipcall (v0.3), adjusted to use modified minimap2 alignment settings accounting for the high variability in the MHC region *(-x asm20 -m 10000 -z 10000,50 -r 50000 --end-bonus=100 --secondary=no --cs -O 5,56 -E 4,1 -B 5*). The reference sequence was hg38 chr6 masked between the HLA-DRA and HLA-DRB1 regions (32,494,000-32,565,000). The validity of the alignment settings was checked by extracting the contig sequences across each of the canonical HLA genes and typing with GenDx (v2.20.2) in PacBio Consensus mode. The resulting variant calls were restricted to SNPs between 29,657,092 - 33,323,016 and merged into a single VCF using vcftools (v0.1.16). Public Dipcall variant calls across the entire genome were downloaded from the HPRC repository. Calls were restricted to SNPs outside of the MHC region and merged into a single VCF. The two sets of variant calls were combined, and non-variant positions were set to homozygous reference if the position was within a region reported as callable by Dipcall. The same filters for encode black list regions, LCR, centromere/telomere, and indels as were used on the clinical samples just with LiftedOver to hg38. Samples were restricted to the African individuals and the two PC outleir were removed (Supplemental Figure 14). The largest population consists of just 23 African samples (after removing two PC outliers, Supplemental Figure 14) and other populations were too small to perform statistical inference for this study. A scan was run filtering on segmental duplications and another without.

A key reason for exploring the Pangenome samples was to further study the SIRP region, which demonstrated surprisingly strong signal in the clinical samples. The *SIRPA* signal was replicated; however, it's paired gene *SIRPB1* likely artifactual and due to platform limitations. The known copy number variation in *SIRPB1* led to strong misleading signals in the clinical samples [8] (Supplemental Figure 5); however, the signals in *SIRPB1* were not replicated using long-read data from the Pangenome (Figure 4). Furthermore, our detailed follow up with samples from both the 17th IHIW and Pangenome highlights the limitations of SNP array and short-read sequencing data and underscores the need for proper characterization of the MHC and the whole genome through new technologies.

*DRB1* is known to have interesting patterns of extended LD, where depending on the *DRB1* allele, a second mutually exclusive DRB gene, either *DRB3*, *DRB4* or *DRB5*, will typically be expressed [9]. The region of DRB1, intron 5, with the strong signal in the 17$^{th}$ IHIW samples It is known to contain an Alu and a LINE, long transposable elements that hinder accurate mapping of shorter sequencing reads. This particular portion of DRB1 is known to have structural variation and disparate repeat elements results in issues when performing multiple sequence alignment and therefore likely causes artifactual LD. These challenges are reconciled when using the Pangenome and consequently the false LD-ABF peak dissipated across the samples.

**Linkage Disequilibrium Approximate Bayes Factor (LD-ABF) using Log-F prior and Data Augmentation**

The model aims to test for increased linkage between the variant of interest and the neighboring variants, where if a region is dense with more polymorphisms and they appear to be in strong LD with the test SNP this may be a strong indication of balancing selection (Figure 1). Phased individual level haplotype data was used to enable the clearest detection that the test variant of interest is in strong linkage with neighboring variants. To test for association between a given variant and neighboring variants first consider just testing the association between haplotypes of one variant versus one other close by neighboring variant. Take test variant $x_i$, $x_i = \{0,1\}$ where 0 corresponds to the major allele and 1 minor allele and $i$ is the index for the individual, and a neighboring variant $y_i = \{0,1\}$, again corresponds to the major or minor allele. A logistic regression model is a natural choice for the binary outcomes.

$$P(\mathrm{y_i} = 1|x_i, \boldsymbol{\beta}) = logit^{-1}(\beta_{0j} + \beta_{1j}x_i)$$

Where $\beta_{1j}$ corresponds to the log odds ratio of observing the alternate allele for neighboring variant $j$ given we observe the alternate allele for the test variant. A standard frequentist approach may run into issues, it is common to see some SNPs in perfect LD or near perfect LD, this creates complete or quasi-complete separation–or rare variants lead to sparsity comparing linkage which can also results in non-identifiability of the model.

A useful set of techniques to overcome problems of separation and sparsity of data are penalization methods, having both frequentist and Bayesian foundations. Penalty functions are used to drive parameter estimates to zero by incurring a cost on including parameters in the model, where they are typically thought of as being effective in high dimensional settings. Greenland and coauthors [10–13] have done extensive work looking at penalized functions or equivalent Bayesian techniques, especially in settings of binary outcomes, and the implications for reducing bias and mean square error (MSE) in settings of separation and sparsity. Greenland proposes using a class of loss functions proportional the information matrix, $r(\beta) = ln(|I(\boldsymbol{\beta})|)^m$. The penalized log likelihood can be written in the form:

$$p(y_i|x_i, \boldsymbol{\beta}) = l(\boldsymbol{\beta}) + \frac{m}{2}\boldsymbol{\beta} - mlog\left(1 + e^{\boldsymbol{\beta}}\right) \quad (1)$$

Where $l(\beta)$ is the standard Bernoulli log likelihood for logistic regression and the other terms penalty terms a function of the $\beta$ parameters being estimated. This has been demonstrated to be proportional to the posterior distribution with $logF(m, m)$ priors, meaning estimating the function of the penalized log likelihood is equivalent to finding the posterior mode. The $logF$ is in the conjugate family for binomial logistic regression, making it a natural choice in such settings. It can be easily seen that at $m = 0$ is equivalent to the maximum likelihood estimate (MLE)–further at $m = 1$ includes Jeffrey's prior in the one parameter model, which was used in this setting based on the recommendations of Greenland and Mansournia [10]. Using establish data augmentation techniques, a computationally efficient and robust way to get estimates of the posterior modes of the coefficients are calculated [14]. Similar to Cauchy (or t-distribution), the $logF$ priros, increase tail weight or skew the prior distribution, where the $logF$ can provide heavier tails than a multivariate normal but lighter tail than the Cauchy.

To test for association between a test SNP and neighboring variant, we use the approximate Bayes factor. The Bayes factor [15] has been used in a variety of settings including in extensive use GWAS [16–19] and the approximate Bayes factor in this setting plugging in equation (1) is defined as:

$$ABF_j = \frac{p(y|M_{1,j})}{p(y|M_{0,j})} = \frac{l(\beta_0, \beta_1) + \frac{m}{2}\beta_0 - mlog\left(1 + e^{\beta_0}\right) + \frac{m}{2}\beta_1 - mlog\left(1 + e^{\beta_1}\right)}{l(\beta_0) + \frac{m}{2}\beta_0 - mlog(1 + e^{\beta_0})} \qquad (2)$$

Comparing the posterior of the intercept only model $M_{0,j} = logit^{-1}(\beta_{0j})$ versus the model with the neighboring variant $M_{1,j} = logit^{-1}(\beta_{0j} + \beta_{1j}x_i)$. If $m$ were to be set to zero, and the data augmentation omitted, that would leave a simple likelihood ratio test, $\frac{l(\beta_0,\beta_1)}{l(\beta_0)}$. To get the test statistics across the entire neighboring region the product of these ABF between the test SNP and each neighboring SNP in a window of a thousand bases (five hundred bases up and downstream) were used, where the log is taken for computational ease. Monomorphic sites are considered to have uninformative ABF of 1. The final statistic is the log product of the ABF across the entire window then divided by the window size, where both taking log are and dividing by the window size are done for interpretability, plugging in equation (2).

$$logABF = \frac{1}{W} log \prod_{j=1}^{W} \left[\frac{P(y|M_{1,j})}{P(y|M_{0,j})}\right] \qquad (3)$$

For a detailed example walking through the calculations and data augmentation techniques for fast Bayesian estimation, see our online resources (https://tris-10.github.io/LD-ABF/documentation/LD_ABF_toyExample).

**Assessing Prediction of Balancing Selection from Evolutionary Simulations**

To test the effectiveness of LD-ABF in detecting balancing selection, a series of forward time simulations were run using using SLiM 3.0 [20,21] and the model was fit and compared with other balancing selection test statistics. Primary focus will be on the first two sets replicating scenarios as described in previous studies [22,23] to demonstrate relative utility of the new method and the last scenarios looks at more recent balancing selection. For each simulation scenario samples of 10,000 were generated across 10 kilobase (Kb) windows, assuming mutation rate and recombination rates of $2.5x10^{-8}$. An ancestral population is simulated for 100,000 generations and then a split occurs, then three different balancing selection scenarios are simulated. For older events the balancing selection mutation is introduced at the time of the split in the center of the 10kb region and then another 250,000 generations are simulated. Then for the younger events the balancing selection mutation is introduced 150,000 generations after the split and then another 100,000 generations are simulated. Last is the new set of simulations looking at more recent balancing selection that has occurred 10,000 generations ago. In all cases, the balancing selection variant is introduced at the center of the 10Kb region. Three different equilibrium frequencies were simulated, {0.25,0.5,0.75}, assuming heterozygous fitness of $1 + hs$ using a selection coefficient *s* of $10^{-2}$ and over dominance coefficient *h* dependent on the desired equilibrium allele frequency corresponding to {-0.5, 100, 1.5}. This can be derived by setting equation 5.11 of Hartl and Clark [24] to zero to solve for the equilibrium frequency for diploid settings of natural selection in population evolutionary inference.An equilibrium frequency of 0.75 indicates the derived allele is under enough positive selection that it becomes more common than the ancestral allele. These simulations therefore also indicate some level of detection of positive selection as well–assuming the evolutionary pressure is not a selective sweep that is strong enough to induce full fixation of the allele. For each of the nine total scenarios (three-time points vs three equilibrium frequencies) two thousand simulations were run along with an additional neutral set where no balancing selection variant was introduced after the split.

The LD-ABF is compared to the HKA statistic [25], Tajima's D [26], Beta Scan $\beta_{2,std}$ [22,23], and $D_{ng}$ statistic [27]. Both HKA and Tajima's D are classic population genetics tests reflecting enrichment in common alleles relative to expectation under a neutral evolving population. Beta Scan looks at a test statistic and compares the weighted regional mutation rate relative to a neutral estimate, while $D_{ng}$ is the sum of the regional binary correlation [28]. All of the statistics appear to perform best for variants of more ancient origin. For example, in the simulations where the balancing selection variant appears 250,000 generations before completion at an equilibrium allele frequency of 50%, the: LD-ABF has an AUC of 98.3%, Tajima's D — 97.7%, $D_{ng}$ — 97.4%, $\beta_{2,std}$ — 96.9%, and HKA — 83.0% (Supplemental Figure 1 and Supplemental Table 1). When the balancing selection variant is more recent in origin, the improvement in predictive performance is higher for LD-ABF relative to the other methods: LD-ABF has an AUC of 94.4% Tajima's D— 93.5%, $D_{ng}$ — 92.4%, $\beta_{2,std}$ — 90.5%, and HKA — 68.4%. Although the classic Tajima's D appears second best in several cases, its performance appears inconsistent. When examining more recent variants at an equilibrium frequency of 25%, LD-ABF has an AUC of 93.4%, where Tajima's D has an AUC of 85.8%, corresponding to an AUC improvement of 7.6% with the new method. Generally, most of the methods (besides HKA) appear to perform well, but LD-ABF appears to be the most robust, consistently predicting signals of balancing selection with either the top or within 0.1% of the top AUC in all simulation scenarios in the replicated set of simulation scenarios.

In the new supplementary set of simulations, looking at more recent balancing selection (Supplemental Figure 1 and Supplemental Table 1), the LD-ABF appears to outperform the $D_{ng}$ more consistently showing up to 4.1% improvement. The Tajima's D outperforms all when the allele frequency is 50% showing 70.9% AUC; however, is again somewhat volatile in it's ability to detect selection showing only 53.6% AUC. All methods have a harder time picking up on selection signal when the balancing selection variant arises more recently, this is to be expected that over a shorter time scale the ability to detect selection signal is more difficult for all methods. As sample sizes increase or the relative selection coefficients are stronger it is expected all methods would also perform better. Generally, the trend appears to show LD-ABF may demonstrate more utility the more recently the variant under selection has arisen in the population. Further, all methods appear to either explicitly or implicitly take local SNP density into account in testing for selection; interestingly, this dependence on SNP density, indicates all tests likely be similarly hindered in real data in settings of limited or missing variants in large part due to platform limitations.

1. Choi, Y., Chan, A. P., Kirkness, E., Telenti, A. & Schork, N. J. Comparison of phasing strategies for whole human genomes. *PLOS Genet.* **14**, e1007308 (2018).
2. Delaneau, O., Zagury, J.-F., Robinson, M. R., Marchini, J. L. & Dermitzakis, E. T. Accurate, scalable and integrative haplotype estimation. *Nat. Commun. 2019 101* **10**, 1–10 (2019).
3. Auton, A. *et al.* A global reference for human genetic variation. *Nature* **526**, 68–74 (2015).
4. Galinsky, K. J. *et al.* Fast Principal-Component Analysis Reveals Convergent Evolution of ADH1B in Europe and East Asia. *Am. J. Hum. Genet.* **98**, 456–472 (2016).
5. Price, A. L. *et al.* Principal components analysis corrects for stratification in genome-wide association studies. *Nat. Genet.* **38**, 904–9 (2006).
6. Amemiya, H. M., Kundaje, A. & Boyle, A. P. The ENCODE Blacklist: Identification of Problematic Regions of the Genome. *Sci. Rep.* **9**, 1–5 (2019).
7. Landrum, M. J. *et al.* ClinVar: Public archive of relationships among sequence variation and human phenotype. *Nucleic Acids Res.* **42**, 980–985 (2014).
8. Royo, J. L. *et al.* A common copy-number variant within SIRPB1 correlates with human out-of-Africa migration after genetic drift correction. *PLoS One* **13**, 1–17 (2018).
9. Kennedy, A. E., Ozbek, U. & Dorak, M. T. What has GWAS done for HLA and disease associations? *Int. J. Immunogenet.* **44**, 195–211 (2017).
10. Greenland, S. & Mansournia, M. A. Penalization, bias reduction, and default priors in logistic and related categorical and survival regressions. *Stat. Med.* **34**, 3133–3143 (2015).
11. Greenland, S. Bayesian perspectives for epidemiological research. II. Regression analysis. *Int. J. Epidemiol.* **36**, 195–202 (2007).
12. Greenland, S. Generalized conjugate priors for Bayesian analysis of risk and survival regressions. *Biometrics* **59**, 92–99 (2003).
13. Mansournia, M. A., Geroldinger, A., Greenland, S. & Heinze, G. Separation in Logistic Regression: Causes, Consequences, and Control. *Am. J. Epidemiol.* **187**, 864–870 (2018).
14. Canario, R. The Best Classifier for Small Datasets : Log- F ( m , m ) Logit. *Medium* 1–15 (2020).
15. Kass, R. E. & Raftery, A. E. Bayes factors. *J. Am. Stat. Assoc.* **90**, 773–795 (1995).
16. Wakefield, J. Bayes factors for genome-wide association studies: comparison with P - values. *Genet. Epidemiol.* **33**, 79–86 (2009).
17. Wakefield, J. A bayesian measure of the probability of false discovery in genetic

epidemiology studies. *Am. J. Hum. Genet.* **81**, 208–227 (2007).

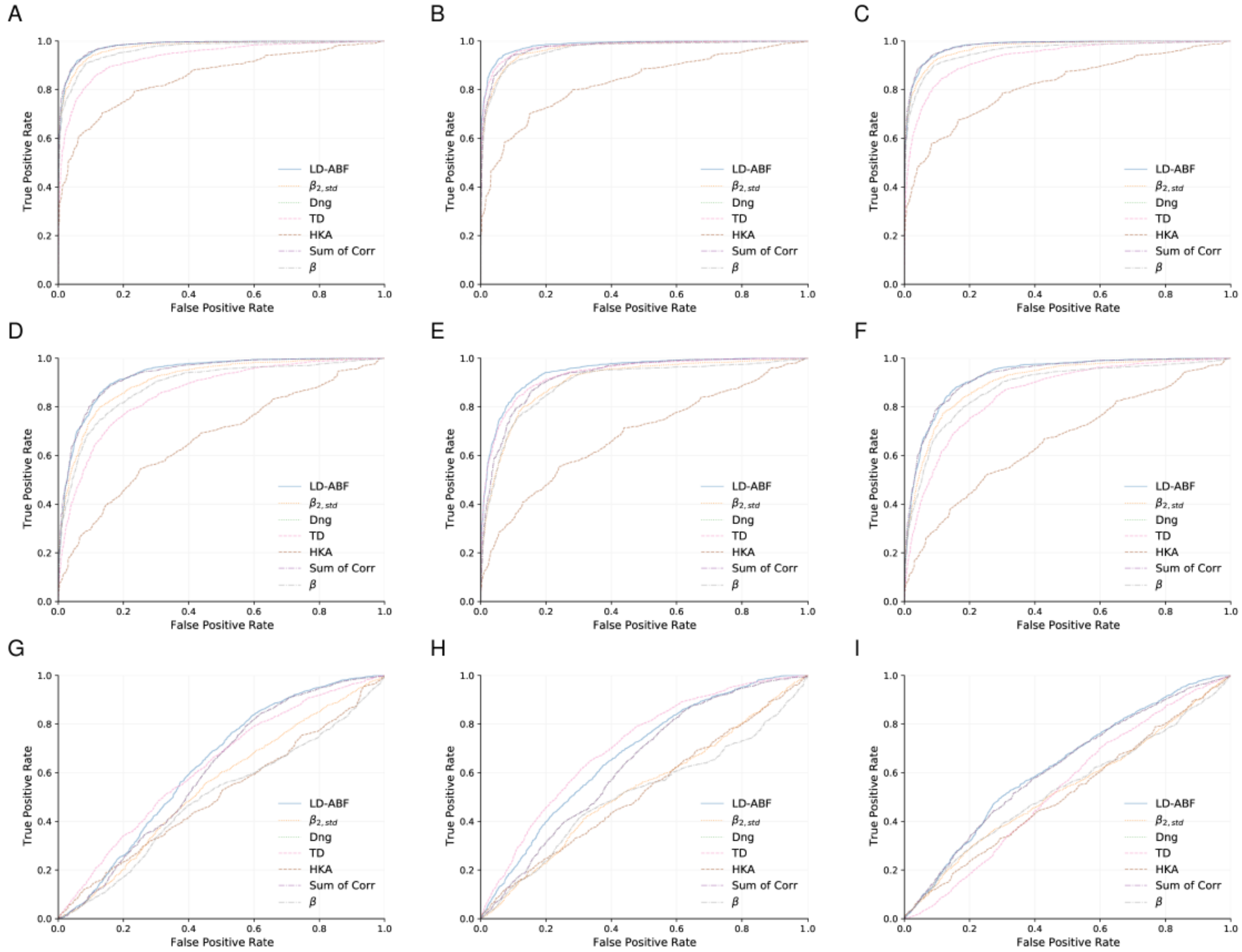


***Supplemental Figure 1 Evolutionary simulation comparison of methods' ability to detect balancing alleles relative to neutral alleles in different scenarios of equilibrium allele frequency and time when mutation is introduced.*** *The top row (A, B, C) correspond to older mutations (250,000 generations before completion) versus the second row (D, E, F) corresponding to younger mutations (100,000 generations back) replicating similar setups described by Stiewart and Voights [29,30] and the bottom row corresponds to recent variants occurring (10,000 generations back). The left column corresponds to a derived allele frequency of 0.25 (A, D, and G), then 0.5 for the middle (B, E, and H), and 0.75 (C, F, and I) for the right column. True positives are taken from the 2,000 balancing selection simulations for each plot and a random common neutral mutation (with MAF>5%) is used from each of the 2,000 neutral simulations to compare as a false positive, ie each plot corresponds to 2,000 simulations half with selection and half without for a given allele frequency and corresponding balancing selection coefficient.*

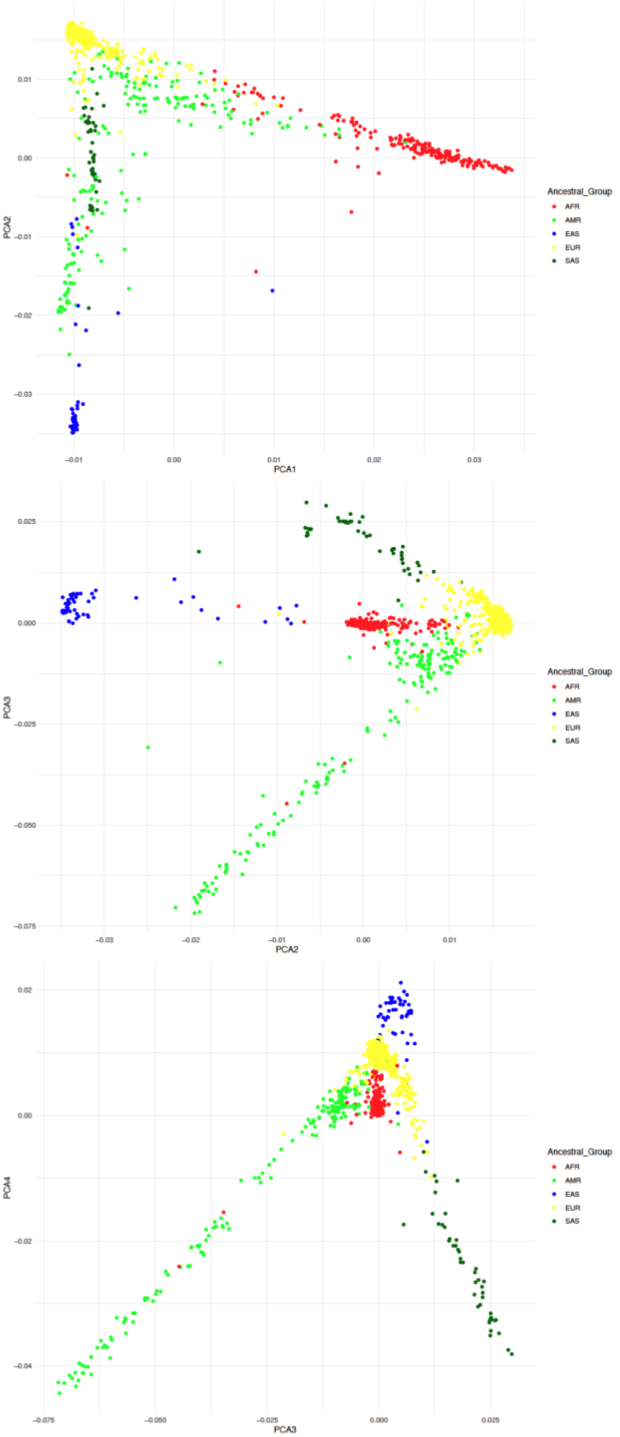


***Supplemental Figure 2 Principal component analysis used for ancestry inference for clinical.*** *Samples are color coded based on closest Thousand Genomes super population they are clustered in where this is the PC analysis before the outliers were removed.*

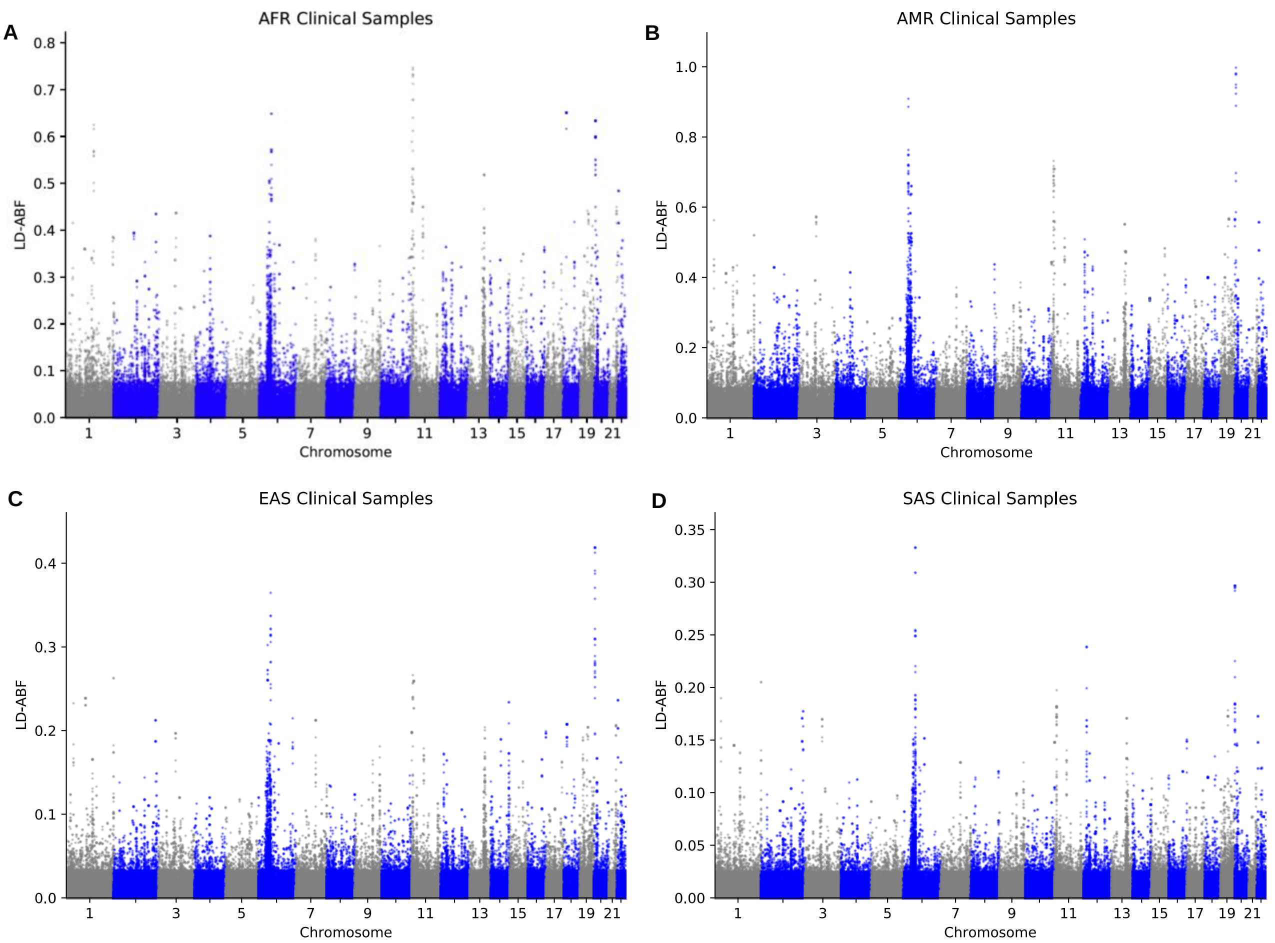


***Supplemental Figure 3 Genome wide scan for balancing selection looking across three different clinical populations: A) AFR B.) AMR C.) EAS and D.) SAS.*** *The relative magnitude of the LD-ABF signals reflect the sample size of the population as any standard test statistic would.*

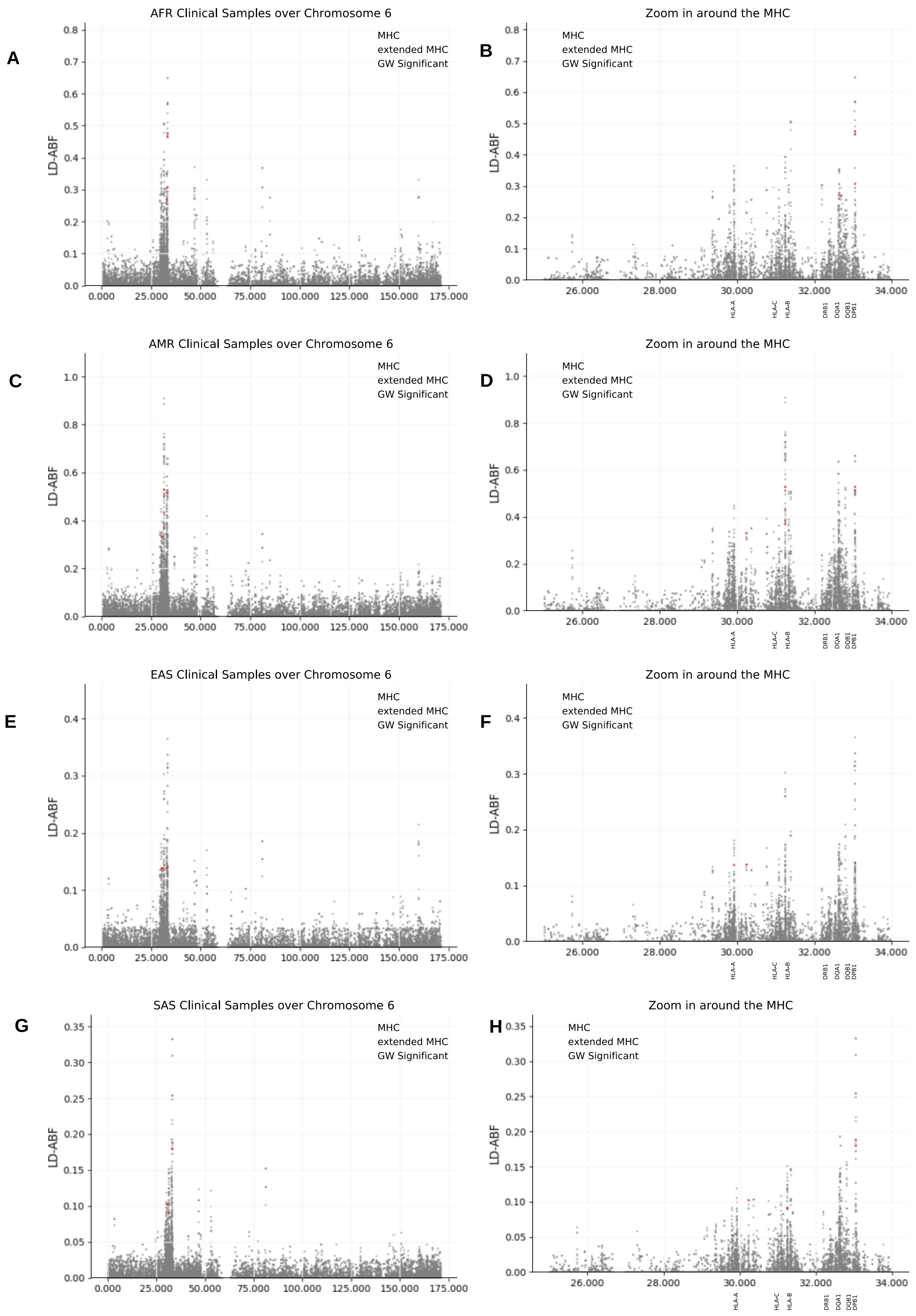


***Supplemental Figure 4 Detailed look at balancing selection chromosome 6 and the MHC in clinical samples.*** *Looking at the A-B) AFR, C-D) AMR, E-F) EAS, and G-H) SAS LD-ABF scan across the clinical samples both over the MHC (A,C,E,G) and zooming in around the MHC region (B,D,E,F).*

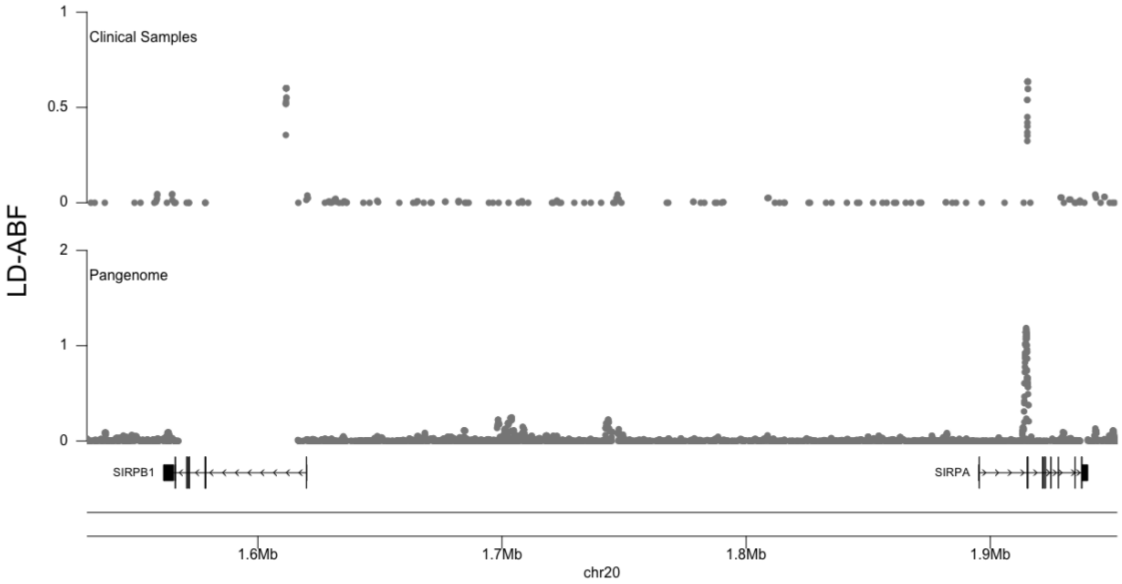


***Supplemental Figure 5 Comparison of LD-ABF over the SIRP region between clinical samples and the Pangenome.*** *The clinical samples are in the top pane which is based off combined exome and SNP array sequencing versus the Pangenome with high quality long-read sequencing. The signal over SIRPB1 does not appear in the Pangenome samples, coupled with the knowledge that the gene has structure variation within it suggesting the signal in the clinical samples is likely artifactual due to mapping or alignment issues.*

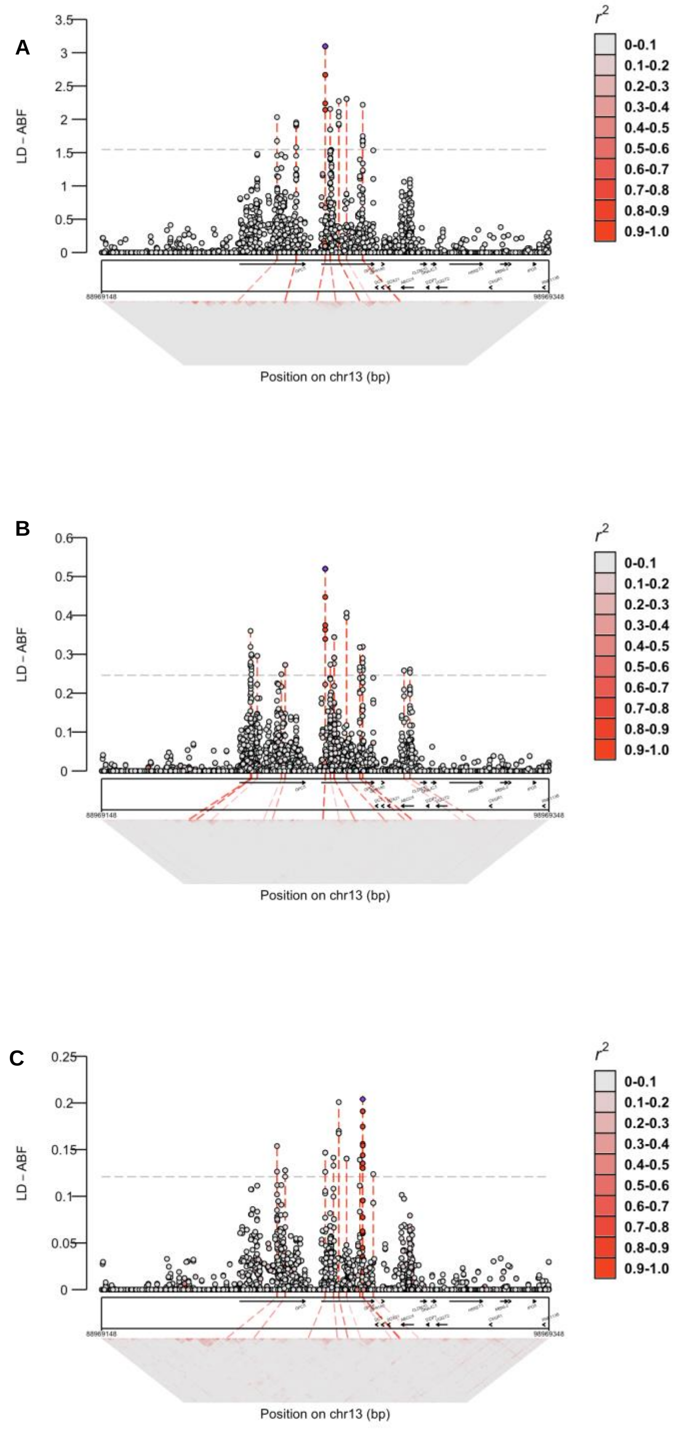


***Supplemental Figure 6** selection signal around GPC genes with LD heat maps below for 3 different CHOP populations: A) EUR B) AFR and C) EAS*

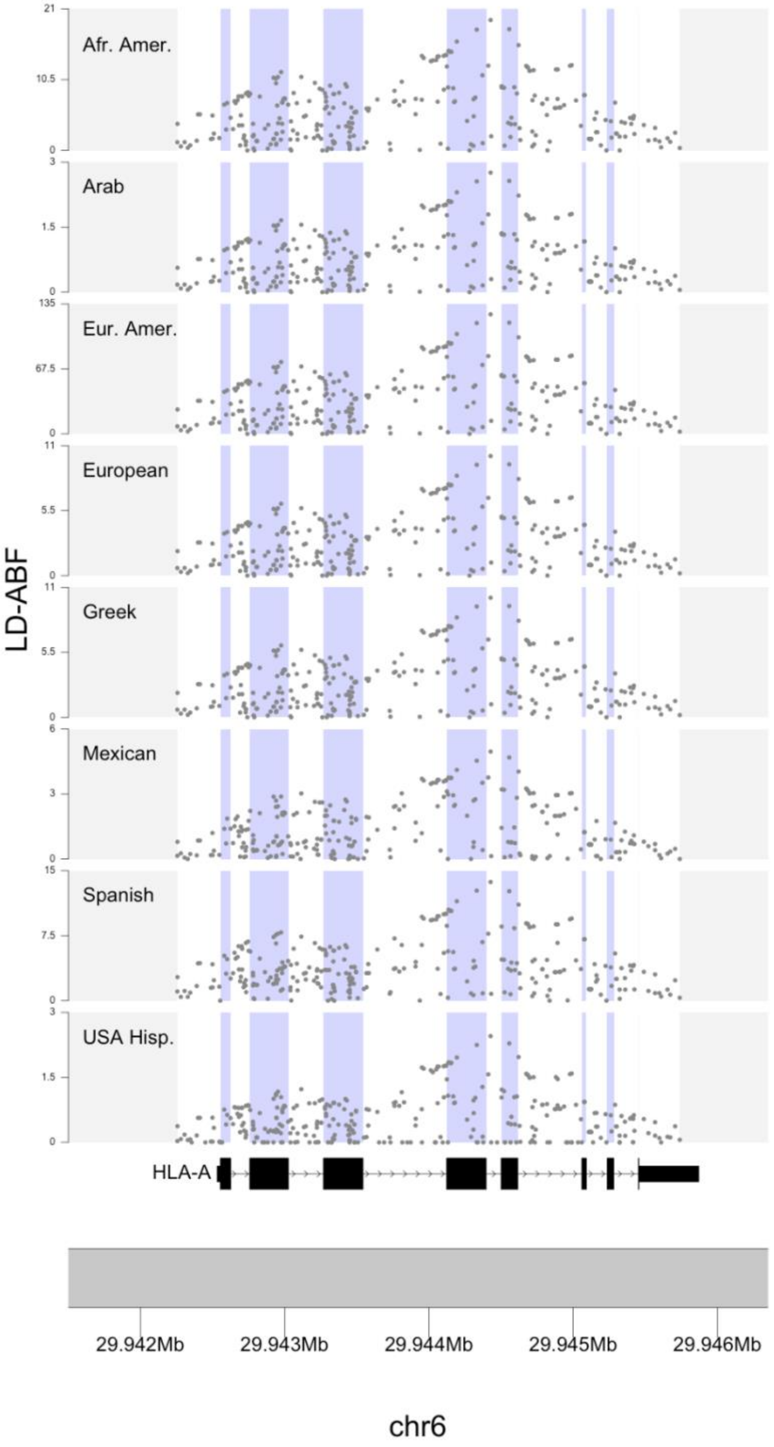


*Supplemental Figure 7 Comparison of LD-ABF across 17th IHIW populatoins for HLA-A*

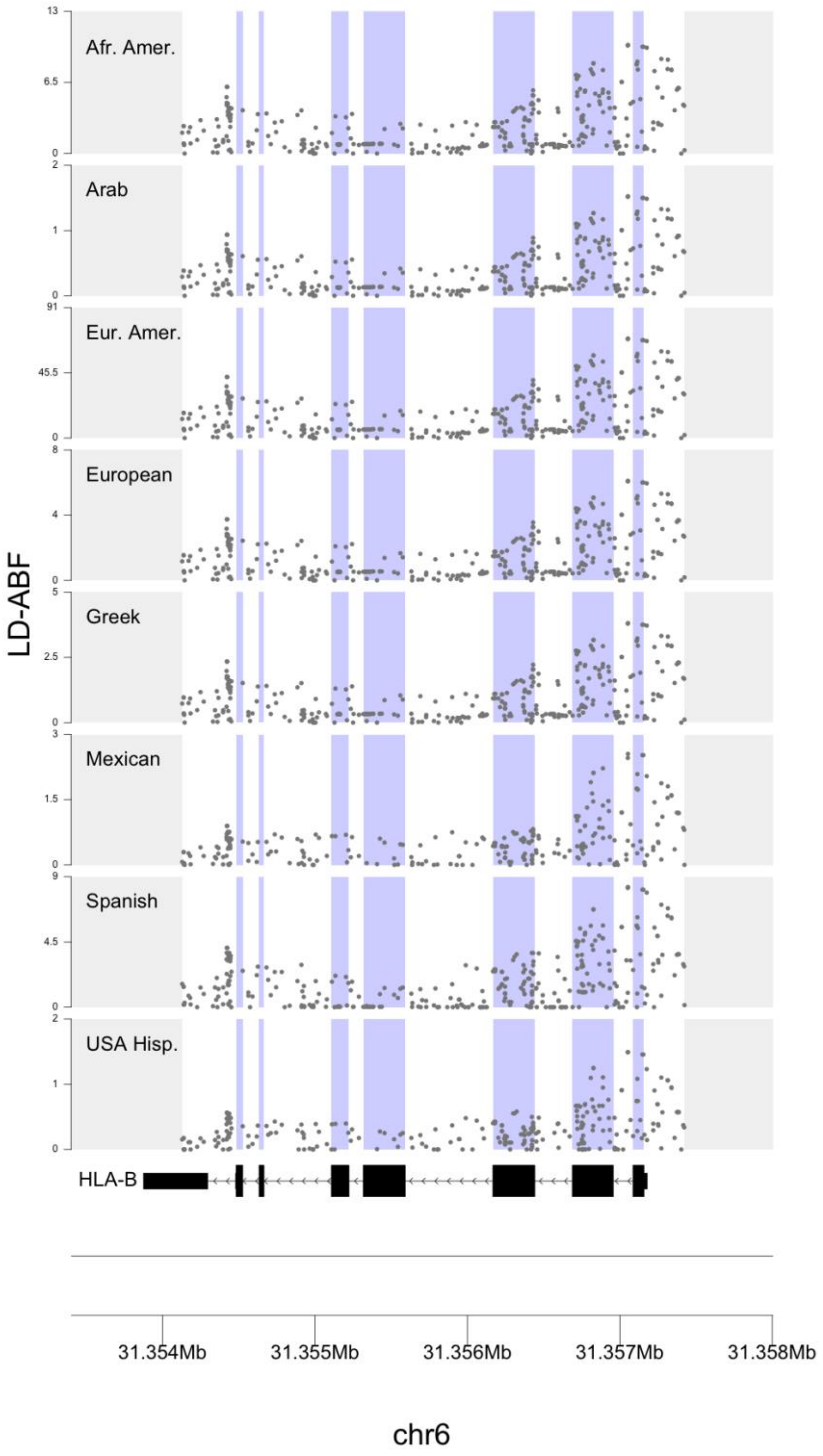


***Supplemental Figure 8 Comparison of LD-ABF across 17th IHIW populations for HLA-B***

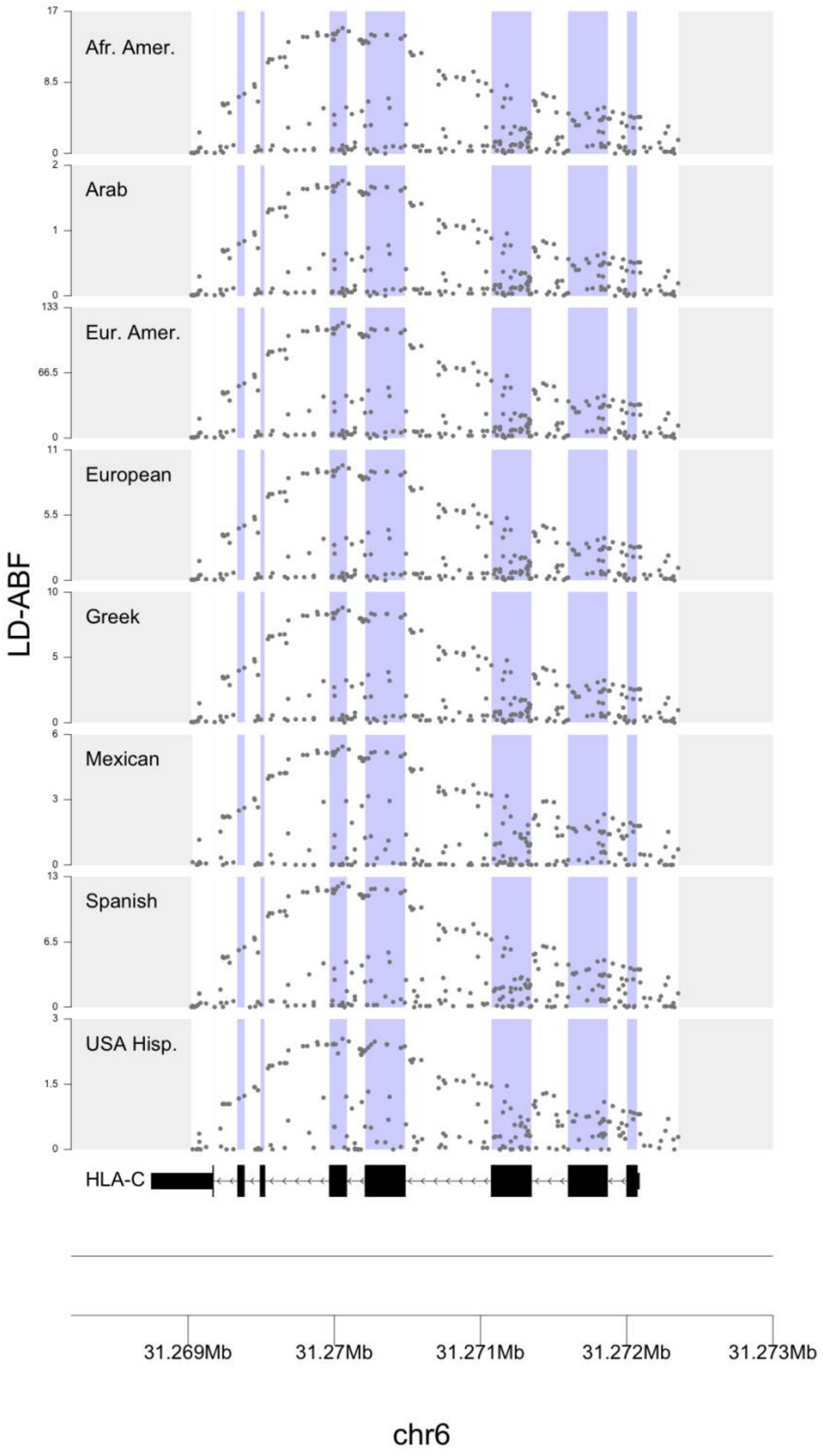


***Supplemental Figure 9 Comparison of LD-ABF across 17th IHIW populations for HLA-C***

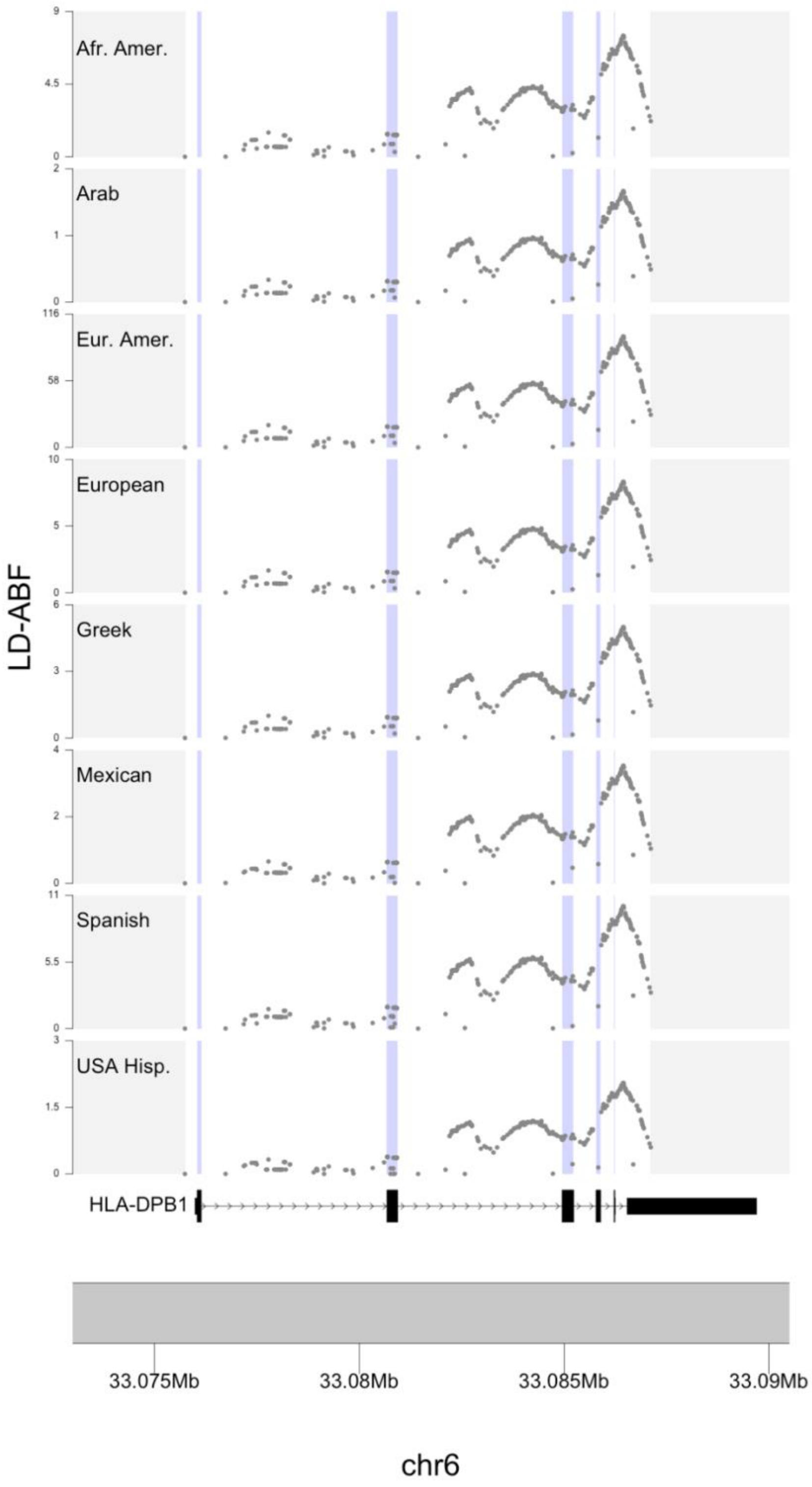


***Supplemental Figure 10 Comparison of LD-ABF across 17th IHIW populations for HLA-DPB1***

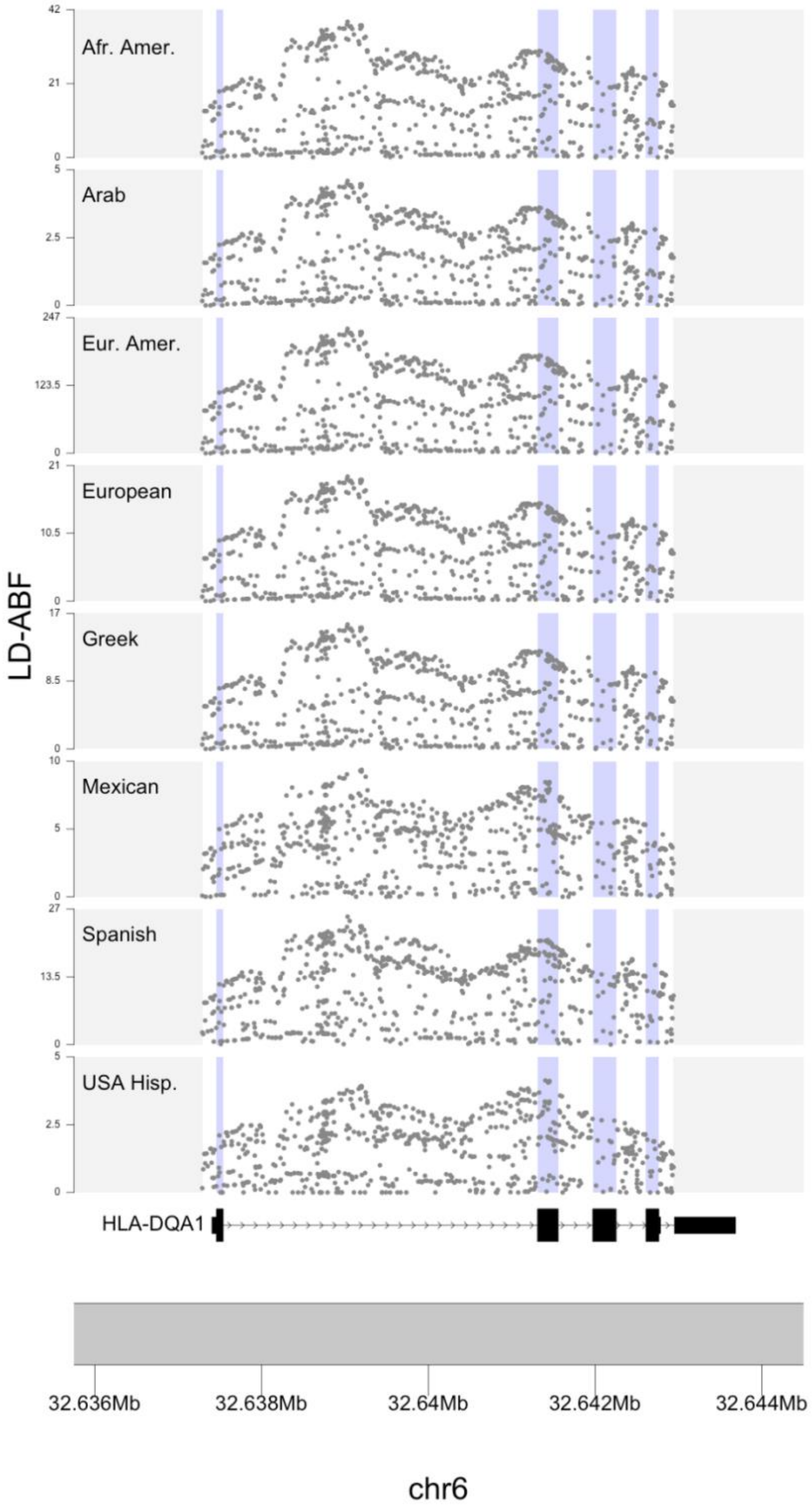


***Supplemental Figure 11 Comparison of LD-ABF across 17th IHIW populations for HLA-DQA1***

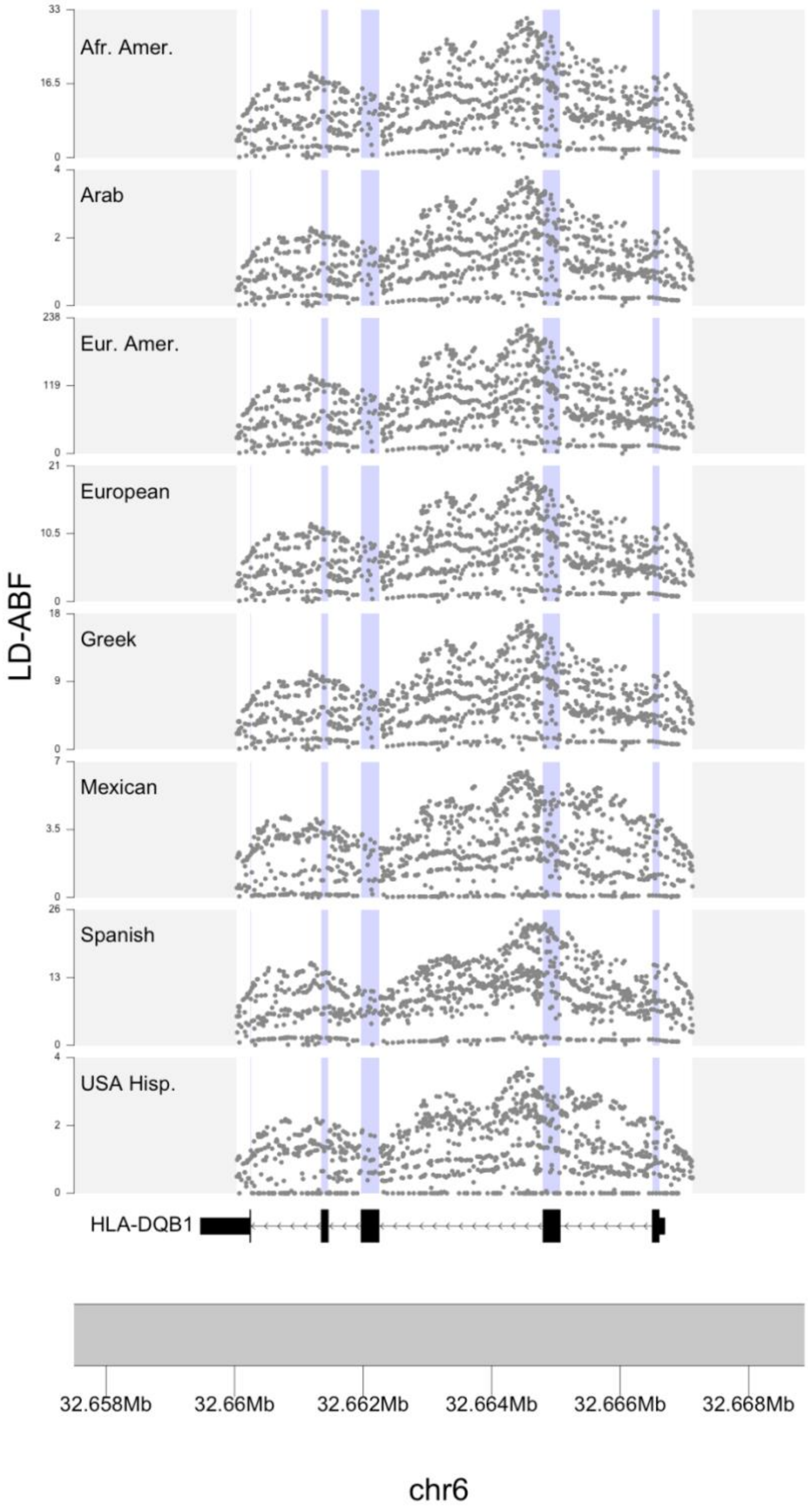


***Supplemental Figure 12 Comparison of LD-ABF across 17th IHIW populations for HLA-DQB1***

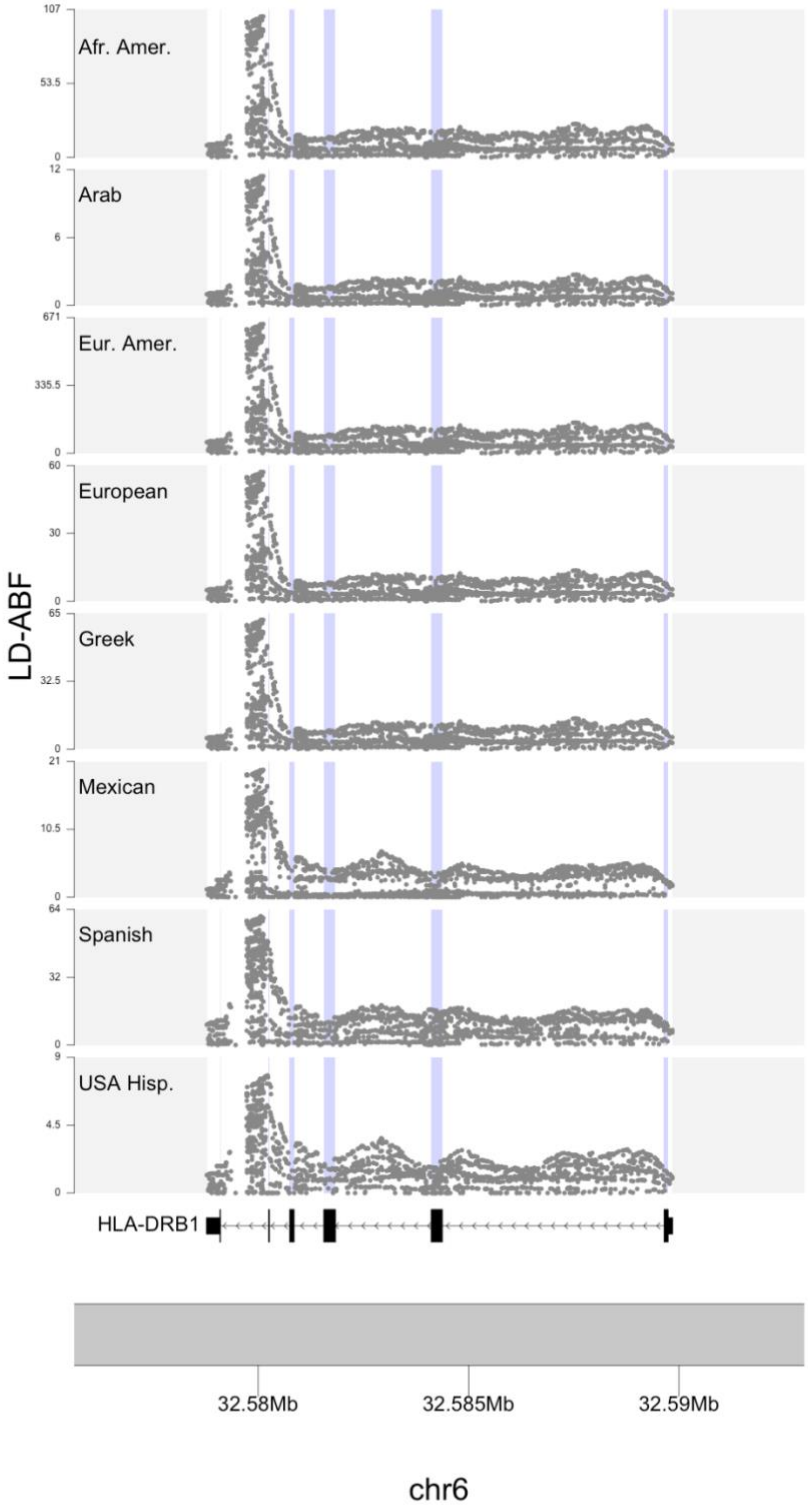


***Supplemental Figure 13 Comparison of LD-ABF across 17th IHIW populations for HLA-DRB1***

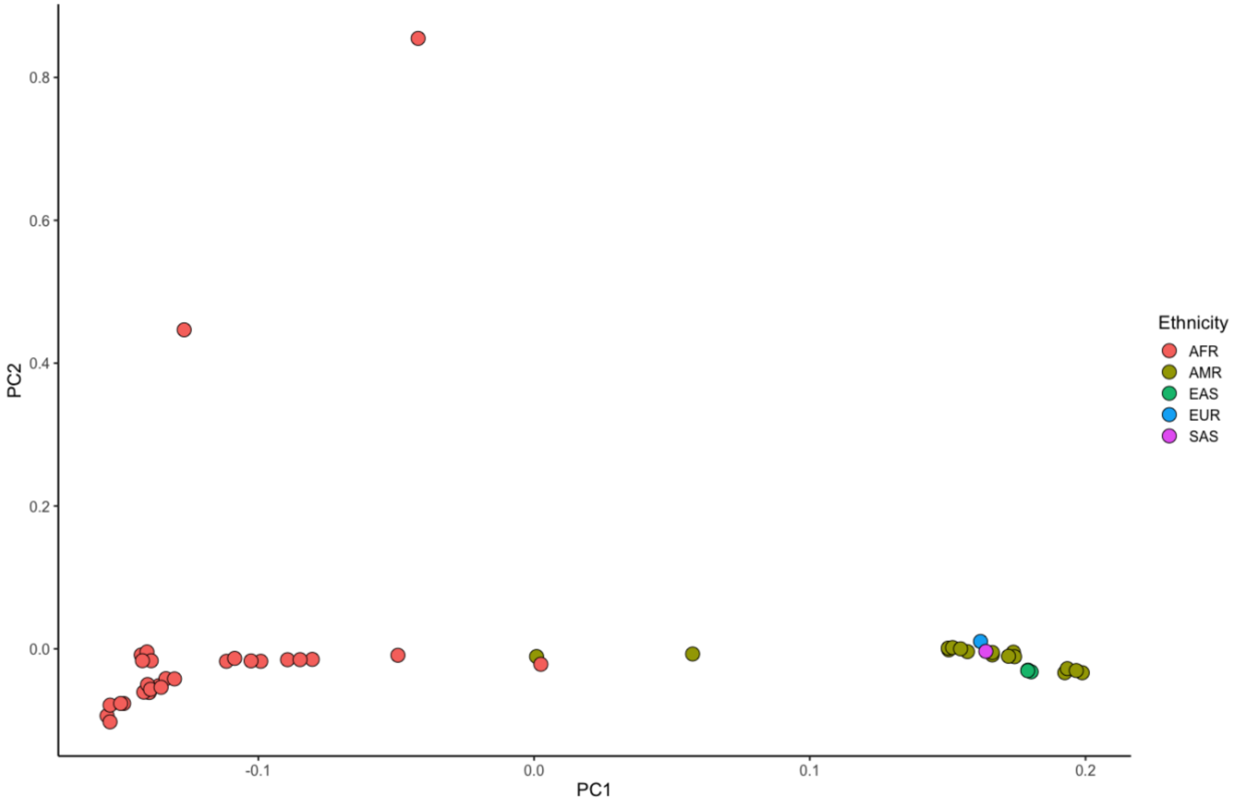


***Supplemental Figure 14 PCA analysis of Pangenome samples including the two outlier African samples before they were removed.***

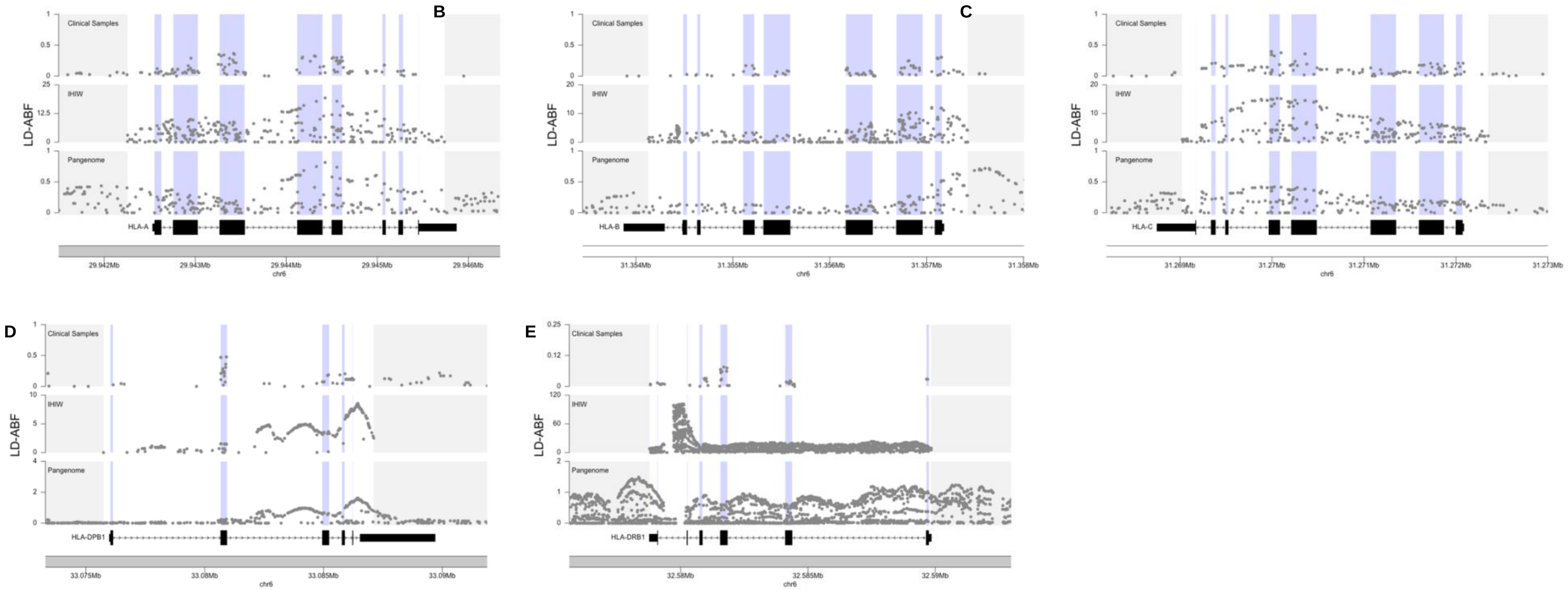


***Supplemental Figure 15* Balancing selection comparison over HLA genes in the the clinical samples, 17th IHIW, and Pangenome**. *Looking in the African American samples within the three different data sets in the 17th IHIW HLA genes A) HLA-A, B) HLA-B, C) HLA-C, D) DPB1, and E) DRB1, while DQA1 and DQB1 can be found in Figure 3.*

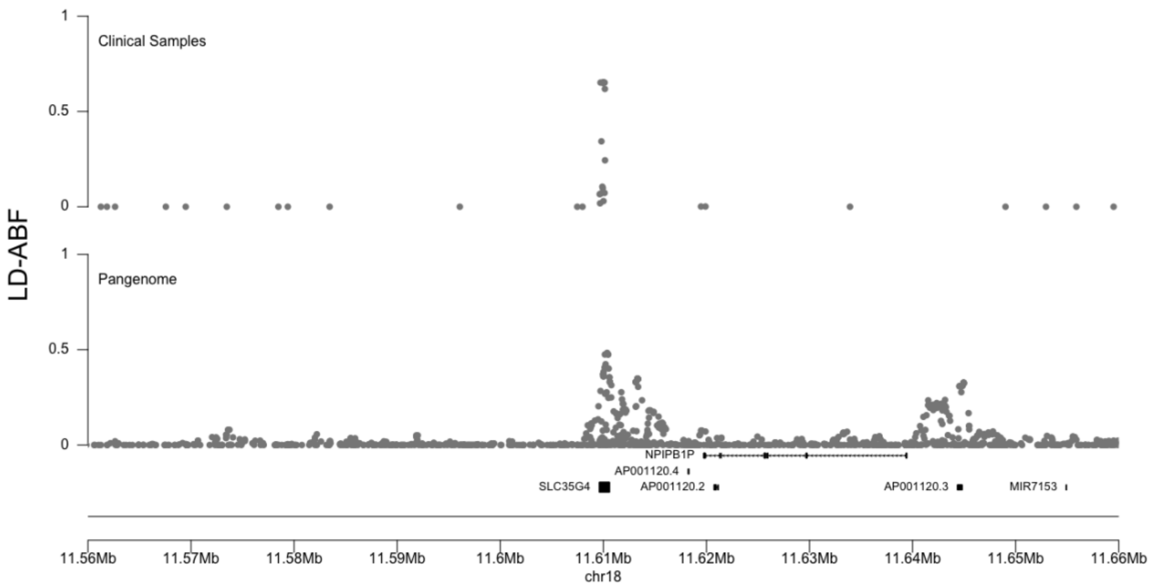


***Supplemental Figure 16 Comparison of LD-ABF around the SLC35G4 in the AFR samples from CHOP trios and the Pangenome.*** *Both samples show a peak in LD density within the solute carrier gene.*

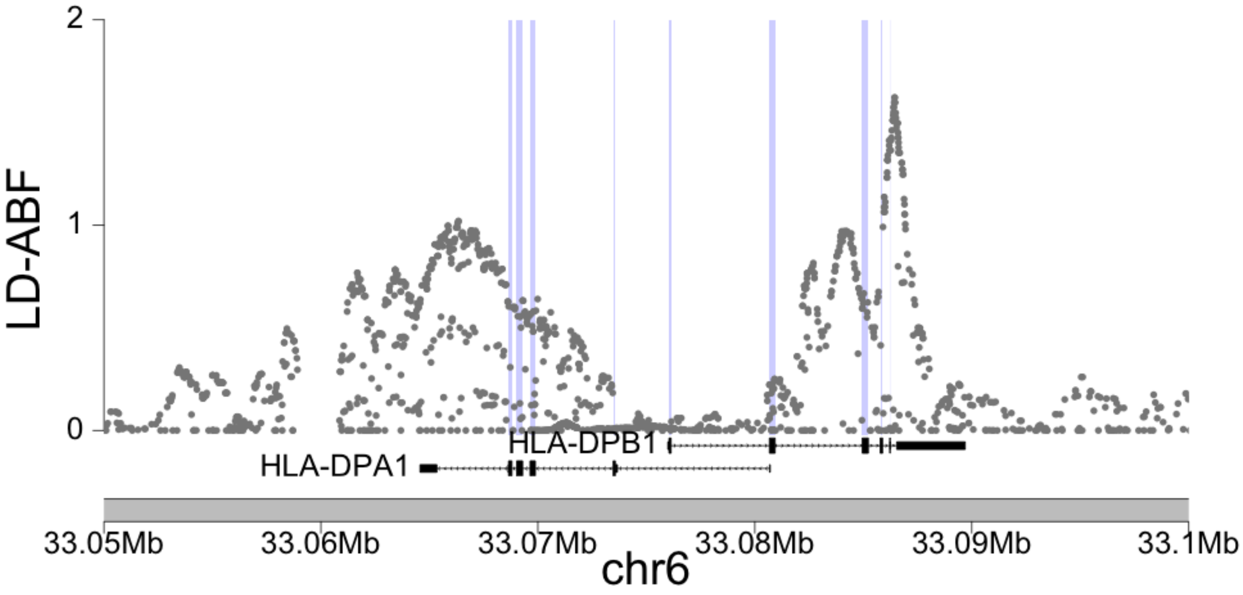


***Supplemental Figure 17 LD-ABF over the DP region in the Pangenome samples.*** *The DPA1 and DPB1 genes overlap and are on reverse strands, where the overlapping sequence of the two genes shows a dip in the relative LD-ABF.*

| *Time* | *h* | *Allele Freq* | *LD-ABF* | $D_{ng}$ | $\beta$ | $\beta_{2,std}$ | *HKA* | *Tajima's D* |
|---|---|---|---|---|---|---|---|---|
| *Old* | -0.5 | 0.25 | 98.1% | 98.2% | 96.4% | 97.5% | 85.0% | 93.5% |
| | 100 | 0.5 | 98.3% | 97.4% | 96.4% | 96.9% | 83.0% | 97.7% |
| | 1.5 | 0.75 | 97.9% | 98.0% | 95.9% | 97.2% | 81.7% | 93.3% |
| Young | -0.5 | 0.25 | 93.4% | 93.3% | 88.7% | 90.9% | 67.1% | 85.8% |
| | 100 | 0.5 | 94.4% | 92.4% | 89.7% | 90.5% | 68.4% | 93.5% |
| | 1.5 | 0.75 | 92.8% | 92.6% | 88.0% | 90.4% | 66.2% | 85.4% |
| Recent | -0.5 | 0.25 | 62.8% | 60.4% | 49.6% | 54.6% | 50.8% | 62.7% |
| | 100 | 0.5 | 67.4% | 63.3% | 51.1% | 53.0% | 52.1% | 70.9% |
| | 1.5 | 0.75 | 62.5% | 61.7% | 53.3% | 53.1% | 51.4% | 53.6% |

***Supplemental Table 1 the AUC for statistics in simulation comparison of methods ability to detect balancing alleles relative to neutral alleles in different scenarios of equilibrium allele frequency and time when mutation is introduced.*** *This corresponds to the same sets of simulations found in Supplemental Figure 1.*

| Pop | CHROM | ID | LD-ABF | GENE(S) | DISEASE/TRAIT | CONTEXT |
|---|---|---|---|---|---|---|
| **EAS** | 20 | rs17855611 | 0.27 | SIRPA | Blood protein levels | Missense |
| | 6 | rs34794906 | 0.17 | HLA-C | Reticulocyte count | Synonymous |
| | 6 | rs1126506 | 0.14 | HLA | Anti-rubella virus IgG levels | Splice region |
| | 6 | rs2858331 | 0.14 | HLA-DQA2 | IgE levels, IgE levels | Regulatory |
| | 6 | rs2516703 | 0.14 | HCG17 | Itch intensity from mosquito bite | Intronic, Intronic |
| | 6 | rs9260151 | 0.14 | MHC, HLA-A | C-peptide levels in type I diabetes | Non coding transcript |
| | 9 | rs8176743 | 0.13 | ABO | End-stage coagulation, Intraocular pressure, Mean corpuscular volume, Mean corpuscular hemoglobin, Blood protein levels | Missense |
| | 9 | rs8176746 | 0.13 | ABO | Mean corpuscular hemoglobin concentration, Mean corpuscular volume, Blood protein levels | Missense |
| | 9 | rs8176749 | 0.13 | ABO | Tumor biomarkers, Urinary metabolites (H-NMR features), Mean corpuscular volume, Venous thromboembolism, Mean corpuscular hemoglobin concentration | Synonymous |
| | 9 | rs8176749 | 0.13 | ABO, Y_RNA, LCN1P2 | Blood protein levels | Synonymous |
| | 6 | rs1050451 | 0.13 | HLA-B, HLA-C | IgG galactosylation phenotypes | Missense |
| | 1 | rs4524 | 0.12 | F5 | Venous thromboembolism | Missense |
| | 1 | rs4525 | 0.12 | F5 | Blood protein levels | Missense |
| | 22 | rs5771225 | 0.12 | SELO | Late-onset Alzheimer's disease | Missense |
| | 9 | rs8176741 | 0.12 | ABO | Elevated serum carcinoembryonic antigen levels, Intraocular pressure | Synonymous |
| | 9 | rs8176741 | 0.12 | RALGDS | Blood protein levels in cardiovascular risk | Synonymous |
| | 9 | rs8176747 | 0.12 | ABO | Platelet count, Reticulocyte count, Blood protein levels, Intraocular pressure | Missense |
| **SAS** | 6 | rs1126506 | 0.19 | HLA | Anti-rubella virus IgG levels | Splice region |
| | 11 | rs5006884 | 0.18 | OR51B5, OR51B6 | Fetal hemoglobin levels | Missense |
| | 6 | rs9277354 | 0.18 | HLA-DPB1 | Antineutrophil cytoplasmic antibody-associated vasculitis | Frameshift |
| | 6 | rs9277356 | 0.18 | HLA-DPB1 | Response to hepatitis B vaccine | Missense |
| | 20 | rs17855611 | 0.16 | SIRPA | Blood protein levels | Missense |
| | 10 | rs2249694 | 0.11 | CYP2E1 | Obesity-related traits | Intronic |
| | 6 | rs2516703 | 0.10 | HCG17 | Itch intensity from mosquito bite | Intronic, Intronic |
| | 9 | rs8176746 | 0.10 | ABO | Mean corpuscular hemoglobin concentration, Mean corpuscular volume, Blood protein levels | Missense |
| | 9 | rs8176749 | 0.10 | ABO | Tumor biomarkers, Urinary metabolites (H-NMR features), Mean corpuscular volume, Venous thromboembolism, Mean corpuscular hemoglobin concentration | Synonymous |
| | 9 | rs8176749 | 0.10 | ABO, Y_RNA, LCN1P2 | Blood protein levels | Synonymous |
| | 9 | rs8176743 | 0.10 | ABO | End-stage coagulation, Intraocular pressure, Mean corpuscular volume, Mean corpuscular hemoglobin, Blood protein levels | Missense |
| | 6 | rs9260151 | 0.10 | MHC, HLA-A | C-peptide levels in type I diabetes | Non coding transcript |
| | 1 | rs4525 | 0.10 | F5 | Blood protein levels | Missense, Missense |
| | 6 | rs2894204 | 0.09 | HLA-C | Waist-hip ratio | Intronic |
| | 9 | rs8176741 | 0.09 | ABO | Elevated serum carcinoembryonic antigen levels, Intraocular pressure | Synonymous |
| | 9 | rs8176741 | 0.09 | RALGDS | Blood protein levels in cardiovascular risk | Synonymous |
| | 9 | rs8176747 | 0.09 | ABO | Platelet count, Reticulocyte count, Blood protein levels, Intraocular pressure | Missense |

***Supplemental Table 2 top Balancing selection GWAS significant SNPs with strong signals of selection in the top 0.1% looking at the CHOP trios for the EAS and SAS populations (a continuation of Table 3).***

| Population | Gene | POS | ID | LD-ABF | DISEASE/TRAIT | CONTEXT | PUBMEDID |
|---|---|---|---|---|---|---|---|
| *European American* | DQA1 | 32606756 | rs9272535 | 198.97 | Red blood cell count | Missense | 27863252 |
| *European American* | DQA1 | 32606756 | rs9272535 | 198.97 | Chronic lymphocytic leukemia | Missense | 21131588 |
| *European American* | DQB1 | 32632659 | rs9274390 | 160.36 | Autism spectrum disorder or schizophrenia | Missense | 28540026 |
| *European American* | DRB1 | 32556601 | rs28724212 | 159.45 | Autism spectrum disorder or schizophrenia | Intronic | 28540026 |
| *European American* | DQB1 | 32628538 | rs201043192 | 133.09 | Lung function (low FEV1 vs high FEV1) | Intronic | 26423011 |
| *European American* | DRB1 | 32550322 | rs9269853 | 128.65 | Alzheimer's disease (late onset) | Intronic | 30617256 |
| *European American* | DRB1 | 32552095 | rs17885382 | 124.85 | Asparaginase hypersensitivity in acute lymphoblastic leukemia | Missense | 25987655 |
| *European American* | DQA1 | 32606878 | rs9272544 | 123.38 | Granulocyte percentage of myeloid white cells | Missense | 27863252 |
| *European American* | DRB1 | 32554129 | rs9270074 | 114.73 | Autism spectrum disorder or schizophrenia | Intronic | 28540026 |
| *European American* | DQA1 | 32608858 | rs4455710 | 110.42 | Squamous cell carcinoma | Intronic | 26829030 |
| *European American* | DQB1 | 32632887 | rs201184533 | 110.13 | Asthma | Intronic | 30929738 |
| *African American* | DQA1 | 32606756 | rs9272535 | 32.30 | Red blood cell count | Missense | 27863252 |
| *African American* | DQA1 | 32606756 | rs9272535 | 32.30 | Chronic lymphocytic leukemia | Missense | 21131588 |
| *African American* | DQB1 | 32632659 | rs9274390 | 25.27 | Autism spectrum disorder or schizophrenia | Missense | 28540026 |
| *African American* | DQB1 | 32628538 | rs201043192 | 19.30 | Lung function (low FEV1 vs high FEV1) | Intronic | 26423011 |
| *African American* | DQB1 | 32632887 | rs201184533 | 19.04 | Asthma | Intronic | 30929738 |
| *African American* | DQB1 | 32632832 | rs9274407 | 16.98 | Drug-induced liver injury (amoxicillin-clavulanate) | Missense | 21570397 |
| *African American* | DRB1 | 32550322 | rs9269853 | 16.96 | Alzheimer's disease (late onset) | Intronic | 30617256 |
| *African American* | DQA1 | 32608858 | rs4455710 | 16.74 | Squamous cell carcinoma | Intronic | 26829030 |
| *African American* | DRB1 | 32554129 | rs9270074 | 16.72 | Autism spectrum disorder or schizophrenia | Intronic | 28540026 |
| *African American* | DRB1 | 32556601 | rs28724212 | 14.76 | Autism spectrum disorder or schizophrenia | Intronic | 28540026 |
| *African American* | DQA1 | 32606878 | rs9272544 | 14.02 | Granulocyte percentage of myeloid white cells | Missense | 27863252 |

***Supplemental Table 3 top balancing selection signals in 17th IHIW samples at GWAS significant SNPs.*** *Signals of balancing selection found in HLA genes within European American and African American populations from 17th IHIW samples that occur at known GWAS significant SNPs.*

| Gene | HGNC ID (gene) | Group |
|---|---|---|
| **ADGRF5** | HGNC:19030 | Immunoglobulin like domain containing, I-set domain containing |
| **ALPK2** | HGNC:20565 | I-set domain containing |
| **CD200R1** | HGNC:24235 | C2-set domain containing |
| **CD276** | HGNC:19137 | C2-set domain containing, V-set domain containing |
| **HLA-A** | HGNC:4931 | C1-set domain containing |
| **HLA-B** | HGNC:4932 | C1-set domain containing |
| **HLA-C** | HGNC:4933 | C1-set domain containing |
| **HLA-DPA1** | HGNC:4938 | C1-set domain containing |
| **HLA-DQA1** | HGNC:4942 | C1-set domain containing |
| **HLA-DQB1** | HGNC:4944 | C1-set domain containing |
| **HLA-DRB5** | HGNC:4953 | C1-set domain containing |
| **HLA-G** | HGNC:4964 | C1-set domain containing |
| **IL1RL1** | HGNC:5998 | I-set domain containing |
| **LILRA1** | HGNC:6602 | Activating leukocyte immunoglobulin like receptors |
| **LILRA6** | HGNC:15495 | Activating leukocyte immunoglobulin like receptors |
| **LILRB2** | HGNC:6606 | Inhibitory leukocyte immunoglobulin like receptors |
| **MICA** | HGNC:7090 | C1-set domain containing |
| **SIGLEC16** | HGNC:24851 | C2-set domain containing, I-set domain containing, Sialic acid binding Ig like lectins, V-set domain containing |
| **SIRPA** | HGNC:9662 | C1-set domain containing, V-set domain containing |

***Supplemental Table 4 HGNC defined immunoglobulin superfamily [31,32] genes that were found in the top 100 balancing selection peaks across the CHOP trio samples****. This includes both the set of genes restricting to a 1Mb window and 100Kb window for defining peaks.*

| Pop | Category | Chr | ID | LD-ABF | Gene | Clinical Disease Name | Sequence Context |
|---|---|---|---|---|---|---|---|
| AFR | Risk factor | 7 | rs10954213 | 0.11 | IRF5 | Systemic lupus erythematosus | 3 prime UTR |
| AMR | Drug response | 22 | rs56011157; rs1081000 | 0.15 | CYP2D6 | Tramadol response | Intronic |
| | Drug response | 22 | rs75276289 | 0.15 | CYP2D6 | Tramadol response | Intronic |
| | Drug response | 22 | rs76312385 | 0.15 | CYP2D6 | Tramadol response | Intronic |
| | Drug response | 22 | rs74644586 | 0.15 | CYP2D6 | Tramadol response | Intronic |
| | Drug response | 22 | rs1080996 | 0.15 | CYP2D6 | Tramadol response | Intronic |
| | Drug response | 22 | rs1080995 | 0.15 | CYP2D6 | Tramadol response | Intronic |
| | Risk factor | 7 | rs10954213 | 0.13 | IRF5 | Systemic lupus erythematosus | 3 prime UTR |
| EAS | Risk factor | 7 | rs10954213 | 0.05 | IRF5 | Systemic lupus erythematosus 0 | 3 prime UTR |
| EUR | Drug response | 22 | rs56011157; rs1081000 | 0.76 | CYP2D6 | Tramadol response | Intronic |
| | Drug response | 22 | rs1080996 | 0.76 | CYP2D6 | Tramadol response | Intronic |
| | Drug response | 22 | rs76312385 | 0.75 | CYP2D6 | Tramadol response | Intronic |
| | Drug response | 22 | rs1080995 | 0.75 | CYP2D6 | Tramadol response | Intronic |
| | Drug response | 22 | rs74644586 | 0.74 | CYP2D6 | Tramadol response | Intronic |
| | Drug response | 22 | rs75276289 | 0.70 | CYP2D6 | Tramadol response | Intronic |
| | Drug response | 6 | rs675026 | 0.68 | OPRM1 | Tramadol response | Synonymous, Intronic |
| | Drug response | 6 | rs562859 | 0.68 | OPRM1 | Tramadol response | Synonymous, Intronic |
| SAS | association | 20 | rs2423326 | 0.05 | HAO1 | Calcium oxalate urolithiasis | Intronic |
| | Drug response | 22 | rs56011157; rs1081000 | 0.04 | CYP2D6 | Tramadol response | Intronic |
| | Drug response | 22 | rs75276289 | 0.04 | CYP2D6 | Tramadol response | Intronic |
| | Drug response | 22 | rs76312385 | 0.04 | CYP2D6 | Tramadol response | Intronic |
| | Drug response | 22 | rs74644586 | 0.04 | CYP2D6 | Tramadol response | Intronic |
| | Drug response | 22 | rs1080996 | 0.04 | CYP2D6 | Tramadol response | Intronic |

***Supplemental Table 5 Top balancing selection signals in clinical samples at known ClinVar variants.*** *Variants in the top 1% of LD-ABF looking at clinical samples with clinical interpretations from ClinVar.*
